# Transcriptional dynamics of nitrogen fixation and senescence in soybean nodules: A dual perspective on host and *Bradyrhizobium* regulation

**DOI:** 10.1101/2025.04.02.646848

**Authors:** Ryan DelPercio, Madison McGregor, Stewart Morley, Nazhin Nikaeen, Blake Meyers, Patricia Baldrich

## Abstract

The Soybean–*Bradyrhizobium* symbiosis enables symbiotic nitrogen fixation (SNF) within root nodules, reducing reliance on synthetic N-fertilizers. However, nitrogen fixation is transient, peaking several weeks after *Bradyrhizobium* colonization and declining as nodules senesce in coordination with host development. To investigate the regulatory mechanisms governing SNF and senescence, we conducted a temporal transcriptomic analysis of soybean nodules inoculated with *Bradyrhizobium diazoefficiens* USDA110. Weekly nodule samples (2–10 weeks post-inoculation, wpi) were analyzed using RNA and small RNA sequencing, while acetylene reduction assays assessed nitrogenase activity from 4 to 7 wpi. We identified three major nodule developmental phases: early development (2–3 wpi), nitrogen fixation (3–8 wpi), and senescence (8–10 wpi). Soybean showed extensive transcriptional reprogramming during senescence, whereas *Bradyrhizobium* underwent major transcriptional shifts early in development before stabilizing during nitrogen fixation. We identified seven soybean genes and several microRNAs as candidate biomarkers of nitrogen fixation, including *lipoxygenases* (*Lox*), suggesting roles for oxylipin metabolism. Soy *hemoglobin-2* (*Hb2*), previously classified as non-symbiotic, was upregulated during senescence, implicating oxidative stress responses within aging nodules. Upregulation of the *Bradyrhizobium paa* operon and *rpoH* during senescence suggested metabolic adaptation for survival beyond symbiosis. Additionally, *Bradyrhizobium NIF* gene expression showed stage-specific regulation, with *nifK* peaking at 2 wpi, *nifD* and *nifA* at 2 and 10 wpi, and *nifH*, *nifW*, and *nifS* at 10 wpi. These findings provide insights into SNF regulation and nodule aging, revealing temporal gene expression patterns that could inform breeding or genetic engineering strategies to enhance nitrogen fixation in soybeans and other legume crops.

## INTRODUCTION

Soybean (*Glycine max*) and *Bradyrhizobium* symbiosis contributes to sustainable agriculture by facilitating **symbiotic nitrogen fixation (SNF)**, a biological process that converts atmospheric nitrogen (N₂) into bioavailable ammonia (NH₃), thereby reducing dependence on synthetic nitrogen fertilizers and enhancing soil fertility. SNF occurs within specialized root nodules through a tightly-regulated symbiotic partnership involving complex signaling, metabolic cooperation, and developmental synchronization between plant and bacterial partners (Udvardi and Poole 2013; Ferguson et al. 2019; Wang et al. 2019; Hoang et al. 2020; Chakraborty et al. 2022).

Advances have been made in understanding nodule initiation and establishment (Stacey et al. 2006; Reyero-Saavedra et al. 2017; Kereszt et al. 2018; Laffont et al. 2019; Kim et al. 2023), however, the molecular mechanisms that orchestrate nitrogen fixation activity and subsequent senescence in both the host plant and rhizobia, are less explored (Franck et al. 2018; Kazmierczak et al. 2020; Zhou et al. 2021). During nodule initiation, soybean roots undergo rapid cellular division and morphological changes in response to rhizobial signaling molecules known as Nod factors, facilitating bacterial invasion and subsequent differentiation into nitrogen-fixing bacteroids enclosed within symbiosomes (Kosslak et al. 1987; Indrasumunar et al. 2011). This early nodule developmental process culminates in active nitrogen fixation, whereby bacterial nitrogenase, encoded by rhizobial *nifH, nifD,* and *nifK* genes, converts atmospheric nitrogen into ammonia (Franck et al. 2018; Jimenez-Vicente et al. 2018; Nguyen et al. 2023). The nitrogen fixation stage is characterized by increasing nitrogenase activity, tightly regulated by the oxygen-buffering capacity of host-leghemoglobin and fueled by plant-derived photosynthates (Kuzma et al. 1993; Kuźma et al. 1999; Smagghe et al. 2009; Singh and Varma 2017; Schwember et al. 2019). However, nitrogen fixation in soybean nodules is inherently transient, typically peaking around flowering to early pod filling (Ciampitti et al. 2021), after which nodules undergo genetically programmed senescence, characterized by structural breakdown, metabolic reprogramming, and gradual termination of SNF (Kazmierczak et al. 2020; Zhou et al. 2021).

Unlike some legumes, such as *Medicago* species, which irreversibly differentiate rhizobia into terminal bacteroids (Mergaert et al. 2003; Haag et al. 2013), soybean maintains U-morphotype bacteroids that retain metabolic flexibility (Duan et al. 2021; Mergaert et al. 2006; Oono et al. 2010). As nodules senesce, these bacteroids continue to undergo transcriptional and metabolic reprogramming, gradually transitioning out of symbiosis with the majority reverting back to a free-living state (Van de Velde et al. 2006; Dupont et al. 2012; Yuan et al. 2021; Müller et al. 2001). This adaptation may enhance rhizobial survival in the soil and improve persistence for future symbiotic interactions.

Nodule organogenesis, referred to as nodulation, is primarily controlled by the host ((Ferguson et al. 2019), and involves coordinated systemic and local signaling pathways. The systemic Autoregulation of Nodulation (AON) pathway governs overall nodule numbers through a root-to-shoot-to-root signaling network (Li et al. 2022b; Gresshoff et al. 2023). Following successful rhizobial infection, the soybean *NINa*-miR172c-*NNC1* (NMN) module triggers the production of Rhizobia-Induced CLE peptides (RICs) at the site of nodulation (Wang et al. 2020, 2019). RICs (RIC1 and RIC2) are then translocated to the shoot, where they activate the Nodule Autoregulation Receptor Kinase (NARK) receptor (Reid et al. 2011; Kereszt et al. 2018). Activated NARK reduces shoot-derived miR2111 levels, removing its suppression of the root-expressed *TOO MUCH LOVE* (*TML*) gene, thereby suppressing further nodulation (Tsikou et al. 2018; Gresshoff et al. 2023; Okuma and Kawaguchi 2021; Zhu et al. 2020). Additionally, local perception of external nitrate is mediated by the transcription factors NIN-Like Proteins 1 and 4 (NLP1 and NLP4), which activate Nitrate-Induced CLE peptides (NICs). These peptides interact locally with the receptor kinase NARK in the root to suppress nodulation under sufficient nitrogen conditions (Reid et al. 2011; Lim et al. 2014; Fu et al. 2024; Li et al. 2022b). While these signaling pathways indirectly regulate nitrogen fixation by controlling nodule number, the local regulation of nitrogen fixation within individual nodules—particularly during peak nitrogen fixation and senescence phases—remains underexplored.

In this study, we conducted comprehensive temporal transcriptional profiling of soybean nodules inoculated with *Bradyrhizobium diazoefficiens* USDA110, spanning critical symbiotic stages from early nodule development to active nitrogen fixation and senescence (2–10 wpi). Using integrated analyses of RNA sequencing, small RNA sequencing, and nitrogen fixation measurements via acetylene reduction assays (ARA) (Montes-Luz et al. 2023), we defined distinct transcriptional patterns corresponding precisely to developmental transitions within nodules: early development (2–3 wpi), nitrogen fixation (3–8 wpi), and senescence (8–10 wpi). We identified a clear decoupling between nitrogenase activity and nodule number and mass, underscoring the need for robust molecular biomarkers to accurately assess nitrogen fixation efficiency. Consequently, our study revealed novel candidate biomarkers and key regulatory factors associated with peak nitrogen fixation, including soybean *Lipoxygenase 1* (*Lox1*, Glyma.07G034800), miR397a/b-5p, and the *Bradyrhizobium* colonization pilus gene *cpaB* (AAV28_RS04130). Additionally, we identified molecular biomarkers associated with nodule aging and senescence, such as soybean *asparagine synthetase-1* (*GS-ASN1,* Glyma.11G170300), miR171p, and the *Bradyrhizobium nifH* gene (AAV28_RS05630), as well as previously uncharacterized transcriptional shifts in *Bradyrhizobium* associated with bacterial adaptation and survival beyond symbiosis.

These findings provide an extended temporal view of the transcriptional dynamics of host and rhizobial genes and miRNAs involved in soybean nodule development. Although the expression of most genes and miRNAs was stage-specific, a subset exhibited biphasic expression patterns, likely participating in multiple developmental stages. This comprehensive analysis deepens our understanding of the complex regulatory interplay between soybean and *Bradyrhizobium*, highlighting promising plant and bacterial targets for optimizing SNF efficiency and nodule longevity. Ultimately, this research advances sustainable agricultural practices and could inform future efforts to engineer nitrogen fixation into non-leguminous crops.

## RESULTS

### Nitrogenase Activity Peaks at 5 wpi and Is Decoupled from Nodule Mass and Number

To investigate symbiotic nitrogen fixation (SNF) dynamics in soybean nodules, we conducted acetylene reduction assays (ARA) on intact nodulated root systems weekly, from 4 to 7 weeks post-inoculation (wpi), a period of active nitrogen fixation **(Figure 1A)**. ARA is a widely used proxy for measuring rhizobial nitrogenase activity, providing an indirect estimate of nitrogen fixation rates in nodules (Hardy et al. 1968; Bergersen 1970; Vessey 2004). Our results revealed that nitrogenase activity peaked at 5 wpi, with high biological variability, showing a significant 2.1-fold increase (p = 0.021) compared to 4 wpi, before gradually declining by 0.83-fold from 5 to 7 wpi (p >0.05) **(Supplemental Table 1)**. These findings indicate that peak nitrogen fixation occurs sometime between 4 and 6 wpi in our study, followed by a gradual decline beginning around 5 to 6 wpi. Consistent with previous studies, we found that nitrogen fixation activity typically decreases over several weeks post-peak, between flowering and pod-filling stages (Franck et al. 2018; Ciampitti et al. 2021). However, due to the increasing size of root systems beyond 7 wpi and the volume constraints of our ARA apparatus, we were unable to assess nitrogenase activity beyond this period.

**Figure 1.**
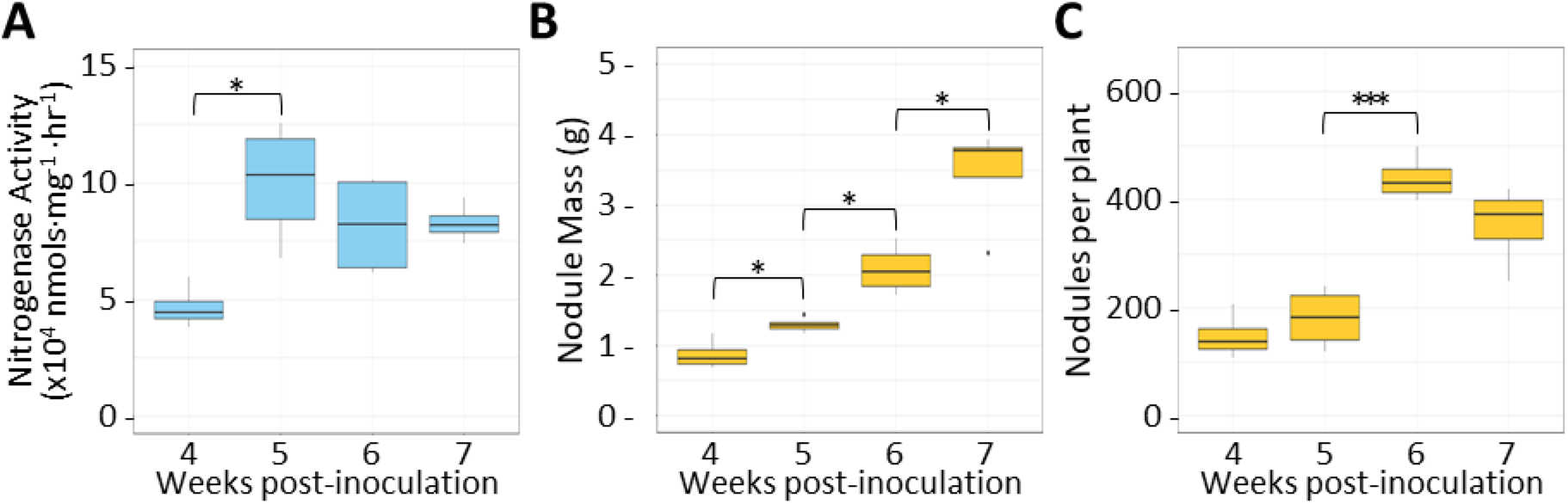
Nitrogenase activity is not contingent upon total nodule mass or number. In the process of symbiotic nitrogen fixation, rhizobial nitrogenase converts atmospheric nitrogen gas into ammonia. Additionally, nitrogenase can fix acetylene gas into ethylene. This ability forms the basis of the acetylene reduction assay (ARA), commonly used to measure nitrogenase activity, an indicator of the level of nitrogen fixation occurring in root nodules. (A) Nitrogenase activity (y-axis) measured via ARA in inoculated soybean whole-root systems (4-7 weeks post inoculation (wpi)) (x-axis). For each timepoint, four experimental root systems and one non-acetylene negative control root system were used. Nitrogenase activity was calculated based on the determined ethylene concentration per time incubated per nodule mass. (B) Fresh weight of nodules collected from whole-plant root systems used in ARA. Nodule mass (y-axis) was measured 3-hours post ARA and used in the calculation of nitrogenase activity. (C) Total number of nodules per plant (y-axis) were counted by hand. Significant p-values are indicated by asterisks: * (0.01 to 0.05), ** (0.001 to 0.01), and *** (< 0.001).

To determine whether nitrogenase activity was associated with nodule traits, we measured nodule mass **(Figure 1B)**, nodule number **(Figures 1C)**, and shoot and root biomass (**Supplemental Figures 1A and 1B**). As expected, all traits increased with plant age. Nodule mass rose significantly each week from 4 to 7 wpi (p < 0.05), but this trend did not mirror the pattern of nitrogenase activity. Similarly, whole-plant nodule numbers increased significantly from 5 to 6 wpi (p = 0.00051), yet this increase was not reflected in nitrogenase levels. While vegetative biomass also increased over time, the substantial rise in nodule mass and number after 5 wpi did not correlate with sustained nitrogenase activity, suggesting that newer nodules contributed minimally to nitrogen fixation. This may result from their developmental immaturity or deregulation of the AON pathway following peak activity or the onset of flowering. In a parallel growth experiment (2–10 wpi), we observed flowering at 5 wpi (6-week-old plants), with visible pod formation by 7 wpi **(Supplemental Figure 2)**. This transition from vegetative to reproductive growth likely plays a central role in regulating the decline in nitrogen fixation, as resources shift toward seed production.

These findings support the idea that host developmental cues govern nodule function and longevity (Ferguson et al. 2019; Gresshoff et al. 2023), a trend further reinforced by our transcriptomic data, and that nitrogenase activity is not directly correlated with nodule number, nodule mass, or host biomass. Given these observations, we next sought to identify candidate molecular biomarkers associated with nitrogen fixation and nodule senescence to improve future assessments of nodulation, including nitrogen fixation activity and symbiotic lifecycle.

### Soybean and *Bradyrhizobium* RNA and sRNA Sequencing Reveal Three Distinct Phases of Nodule Development

To identify coding genes and small RNAs (sRNA) involved in nitrogen fixation and nodule senescence, we examined transcriptomic changes throughout soybean nodule development. We performed dual RNA and small RNA sequencing (RNAseq and sRNA-seq) of soybean nodules formed post inoculation with *Bradyrhizobium diazoefficiens* USDA110, collected weekly from 2 to 10 wpi **(Figure 2)**. Transcriptome mapping revealed that the majority of reads (∼75–80%) consistently aligned to the soybean genome across all time points, while ∼5–10% mapped to *Bradyrhizobium* **(Figure 2A; Supplemental Tables 2 and 3)**. These proportions remained stable throughout nodule development, underscoring the robustness of our dataset for downstream analyses.

**Figure 2.**
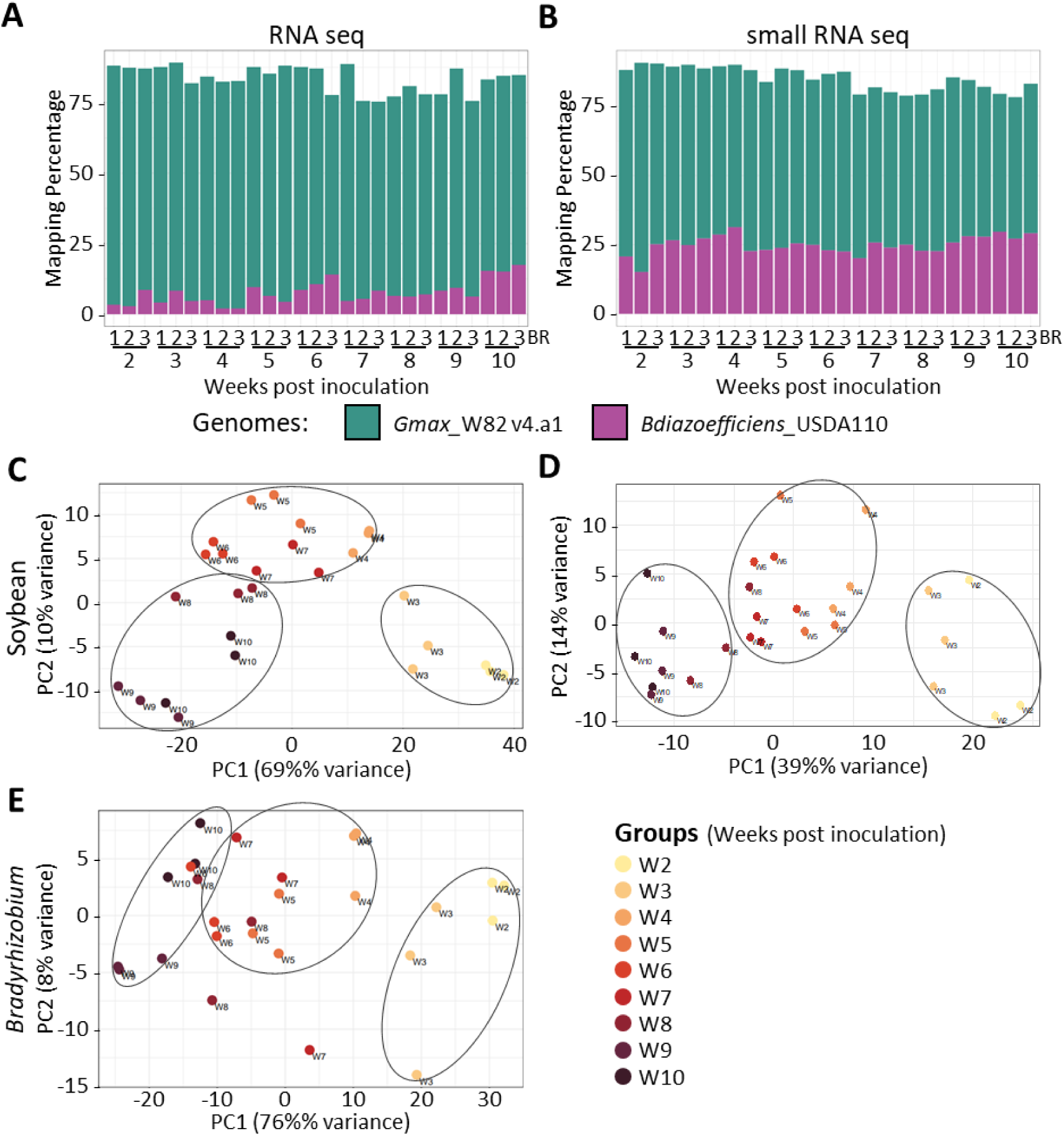
Transcriptomic profiling reveals three distinct stages of soybean nodule development (2-10 weeks post-inoculation) (A, B) Mapping percentages of RNA-seq (A) and small RNA-seq (B) reads to the *Glycine max cv.* Williams-82 and *Bradyrhizobium diazoefficiens* USDA110 genomes across nodule development (2–10 wpi). The majority of reads aligned to the soybean genome (∼75–80% for RNA-seq, ∼60% for sRNA-seq), while *Bradyrhizobium* reads accounted for ∼5–10% of RNA-seq and ∼25% of sRNA-seq reads. (C, D) Principal Component Analysis (PCA) of soybean RNA-seq (C) and small RNA-seq (D) data revealed three major developmental clusters: early development (2–3 wpi), nitrogen fixation (3–8 wpi), and senescence (8–10 wpi). (E) PCA of *Bradyrhizobium* transcriptomic data identified similar phases, with greater variation during early nodule development and pre-peak nitrogen fixation (2–5 wpi). The reduced separation of *Bradyrhizobium* samples in PC2 suggests that host transcriptional shifts drive major symbiotic transitions. These findings highlight host-dominant regulation of nodule development and nitrogen fixation.

Similarly, sRNA-seq confirmed the presence of reads from both plant and bacterial origins, with ∼60% of reads mapping to the soybean genome and ∼25% to *Bradyrhizobium*, while the rest remained unmapped **(Figure 2B; Supplemental Table 4)**. Analysis of sRNA size distributions revealed distinct profiles of plant and bacterial origin. Consistent with known plant sRNA classes (Zhan and Meyers 2023; Borges and Martienssen 2015), soybean-derived sRNAs exhibited three major peaks at 21, 22, and 24 nucleotides **(Supplemental Figure 3A)**. Notably, a shift from a predominant 22-nucleotide sRNA peak during early nodule development (2 wpi) to a dominant 21-nucleotide peak from 4 wpi onward suggests a temporal shift in miRNA processing and regulatory activity during nodule maturation. This observation aligns with known roles of 21-nt and 22-nt miRNAs in triggering transcript cleavage versus secondary siRNA production, respectively (Zhan and Meyers 2023; Chen et al. 2010; Cuperus et al. 2011). In contrast, *Bradyrhizobium* sRNAs exhibited a broader size distribution (17–35 nt) with two dominant peaks at 28 and 30 nt emerging from 7 wpi onward. While the precise functions of these bacterial sRNAs remain uncharacterized, their dynamic accumulation pattern during senescence suggests potential regulatory involvement in *Bradyrhizobium* gene expression during symbiotic decline **(Supplemental Figure 3B)**.

Principal Component Analysis (PCA) of RNA and sRNA expression profiles revealed distinct transcriptional shifts in both soybean and *Bradyrhizobium*, reflecting functional transitions throughout nodule development. Soybean transcriptomic data **(Figure 2C)** clustered into three groups: 2 to 3 wpi, 4 to 8 wpi, and 9 to 10 wpi. Together with our ARA and physiological data, these clusters likely correspond to early development (2–3 wpi), nitrogen fixation (3–8 wpi), and senescence (8–10 wpi). Similarly, plant miRNA accumulation patterns followed these transitions, supporting the hypothesis that miRNAs serve as key regulators of nodule development in coordination with host developmental stage shifts (Hoang et al. 2020; Simon et al. 2009; Yun et al. 2023) **(Figure 2D)**. *Bradyrhizobium* transcriptomic profiles also separated into three phases (2–3 wpi, 4–7 wpi, and 8–10 wpi) with wider variation in PC1, between pre-peak nitrogen fixation (2–5 wpi) than post-peak nitrogen fixation (5–10 wpi), suggesting that *Bradyrhizobium* undergoes greater transcriptional changes during early development and pre-peak nitrogen fixation **(Figure 2E)**. Additionally, we observed less separation between samples in PC2 component in *Bradyrhizobium* compared to soybean, suggesting reduced variation in host between different stages of development, reinforcing the hypothesis that “the host controls the party” (Ferguson et al. 2019).

### *Bradyrhizobium* exhibits early transcriptomic shifts, while soybean undergoes extensive senescence-driven gene regulation

To further characterize these transitions, we performed differential expression (DE) analysis of coding genes and differential accumulation (DA) analysis of sRNAs across weekly comparisons. We identified 7,510 soybean genes that were DE. Of these, 14% (n = 1,025) were DE during early development (2–3 wpi comparison) **(Figure 3A; Supplemental Table 5)**. During nitrogen fixation (3–8 wpi comparisons), the number of DE genes per weekly comparison decreased, suggesting a period of relative transcriptomic homeostasis. Conversely, senescence (8–10 wpi comparisons) exhibited the highest number of DE genes per weekly comparison, accounting for 52% (n = 3,923) of total DE genes, suggesting a period of high transcriptomic regulation in nodules.

**Figure 3.**
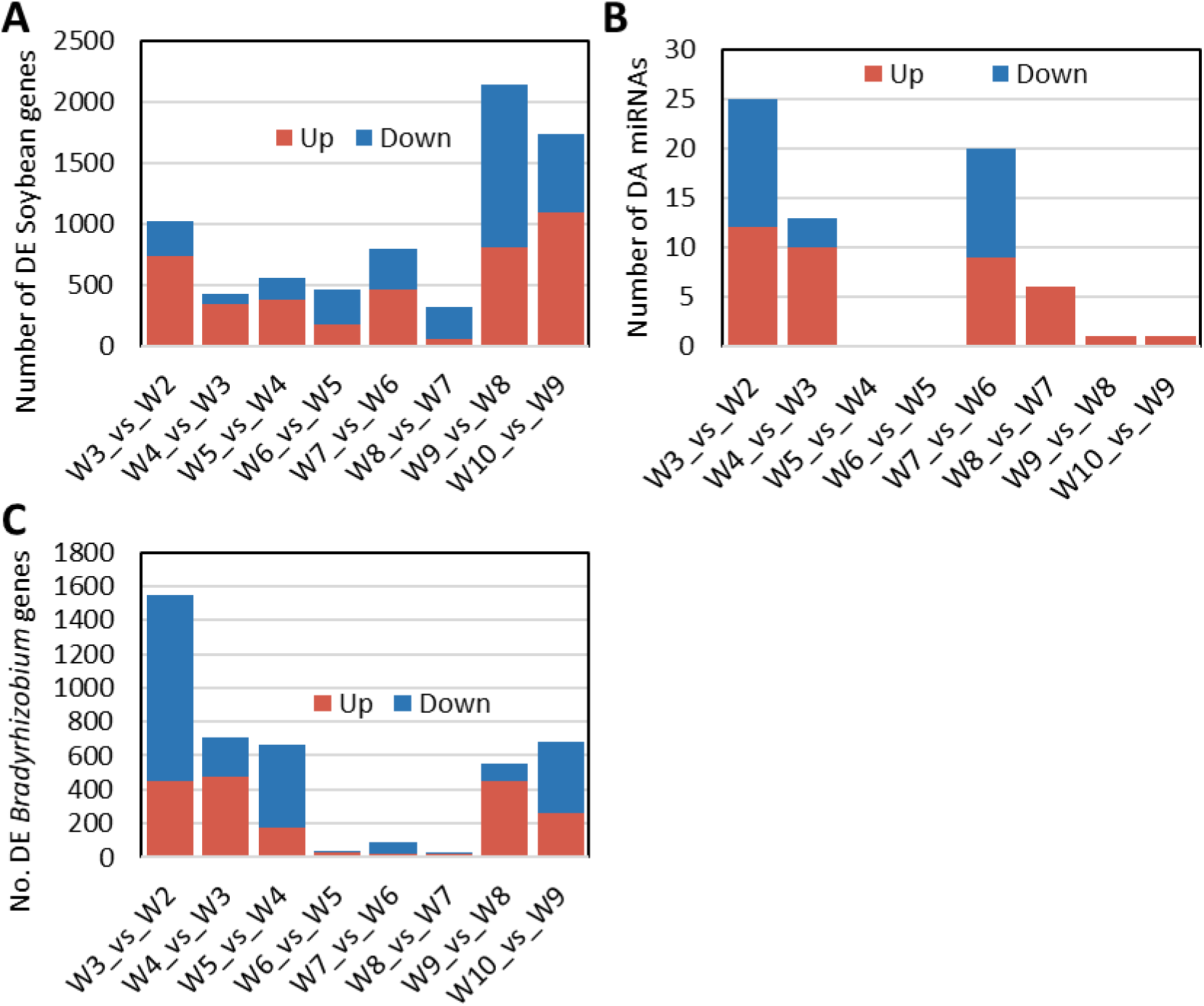
Distinct temporal patterns of differential gene expression in soybean and *Bradyrhizobium* during early nodule development and senescence. Stacked bar plots illustrate the distribution of differentially expressed (DE) genes and differentially accumulated (DA) miRNAs across weekly comparisons (2–10 wpi) in soybean nodules. Upregulated genes and miRNAs shown in red, while downregulated genes and miRNAs shown in blue. (A) Soybean DE genes. Senescence (8–10 wpi) exhibits the highest number of DE genes, while nitrogen fixation (3–8 wpi) is marked by relative transcriptomic stability. (B) Soybean DA miRNAs. Most DA miRNAs accumulate during early development (2–3 wpi) and early senescence (6–7 wpi), suggesting a role in regulating developmental transitions. No miRNAs were DA during peak nitrogen fixation (4–6 wpi). (C) *Bradyrhizobium* DE genes. The highest number of DE genes occurs during early nodule development (2–5 wpi), followed by a sharp decline during nitrogen fixation (5–8 wpi). Expression increases again during senescence (8–10 wpi), suggesting bacterial adaptation to declining nodule conditions.

During early development (2–3 wpi), soybean DE genes were enriched in carbohydrate metabolism, transmembrane transport, transcriptional regulation, and ribosomal biosynthesis, supporting nodule organogenesis and rhizobial infection **(Supplemental Figure 4)**. As nodules transitioned into the nitrogen fixation phase (3–8 wpi), upregulated genes were associated with microtubule-based processes, polysaccharide metabolism, proteolysis, and sulfate transport, reflecting cellular remodeling and metabolic adjustments necessary for SNF. Photosynthetic electron transport and sulfur metabolism were also enriched, underscoring the plant’s coordination of energy and nutrient supply to sustain nitrogen fixation. During senescence (8–10 wpi), DE genes were primarily enriched in nutrient recycling, cell wall degradation, and programmed cell death, while pathways related to methionine biosynthesis, DNA replication, and oxidative stress responses were significantly downregulated, marking the decline of SNF and the functional termination of nodules.

We identified 64 DA miRNAs across weekly comparisons throughout nodule development (2–10 wpi), with accumulation patterns aligning closely to key developmental transitions **(Figure 3B; Supplemental Table 6)**. DA miRNAs were predominantly associated with early development (2–3 wpi), the onset of nitrogen fixation (3–4 wpi), and the early stages of programmed senescence and decline of nitrogen fixation (6–8 wpi). Notably, the majority of DA miRNAs, ∼58% (37/64), were DA prior to peak nitrogen fixation (2–4 wpi), while another ∼41% (26/64) exhibited DA immediately following peak activity (6–8 wpi). In contrast, no miRNAs were DA during a period of active nitrogen fixation (4–6 wpi), and only two were DA during senescence (8–10 wpi). These results suggest that miRNA regulation plays a crucial role in orchestrating developmental phase transitions rather than maintaining nitrogen fixation itself. The enrichment of DA miRNAs prior to nitrogen fixation and again during early senescence suggests that miRNAs predominantly regulate the establishment and termination of symbiosis, facilitating key transcriptomic shifts as nodules transition between functional states.

For *Bradyrhizobium*, we identified a total of 4,299 DE genes across weekly comparisons. We observed the most prominent number of DE genes, ∼68% (2,919/4,299), before peak nitrogen fixation (2–5 wpi). The 2 to 3 wpi comparison comprised more than a third, or 36% (1,551/4,299) of total DE genes, most of which, ∼71% (n=1,101/1,551), were downregulated, corresponding to active bacterial adaptation to the nodule environment **(Figure 3C; Supplemental Table 7)**. Notably, we observed a minimal number, or ∼3% (141/4,299) of DE genes from 5 to 8 wpi, suggesting a homeostatic state in which *Bradyrhizobium* transcriptional regulation remained stable and minimal transcriptional changes are required, similar to soybean DE from 3 to 8 wpi. Conversely, during senescence (8–10 wpi), the number of DE genes per weekly comparison increased again, 29% (1,239/4,299), suggesting an increased bacterial response to deteriorating nodule conditions and potential reprogramming for survival outside the symbiotic relationship.

### Transcriptional regulation is stage-specific, with few genes exhibiting multiple DE events across developmental stages

Building on our transcriptomic analysis, we examined whether DE genes and DA miRNAs were stage-specific or regulated across multiple developmental transitions (multi-stage). We grouped DE genes and DA miRNAs into the three previously described developmental stages. We identified a total of 4,961 soybean and 2,962 *Bradyrhizobium* DE genes, as well as 55 DA miRNAs representing 36 unique mature sequences **(Figure 4)**. These numbers reflect the total number of distinct genes and miRNAs that exhibited one or more instances of differential regulation. We observed that the majority of DE genes were stage-specific, encompassing approximately 80% (3,951 out of 4,961) of soybean DE genes **(Figure 4A; Supplemental Table 8)** and 74% (2,188 out of 2,962) of *Bradyrhizobium* DE genes **(Figure 4B; Supplemental Table 9)**. Likewise, nearly all DA miRNAs, ∼93% (51 out of 55) were stage-specific **(Figure 4C; Supplemental Table 10)**. These findings suggest that transcriptional regulation in both soybean and *Bradyrhizobium* is largely confined to specific developmental stages, though a subset of genes and miRNAs undergo regulation across multiple transitions.

**Figure 4.**
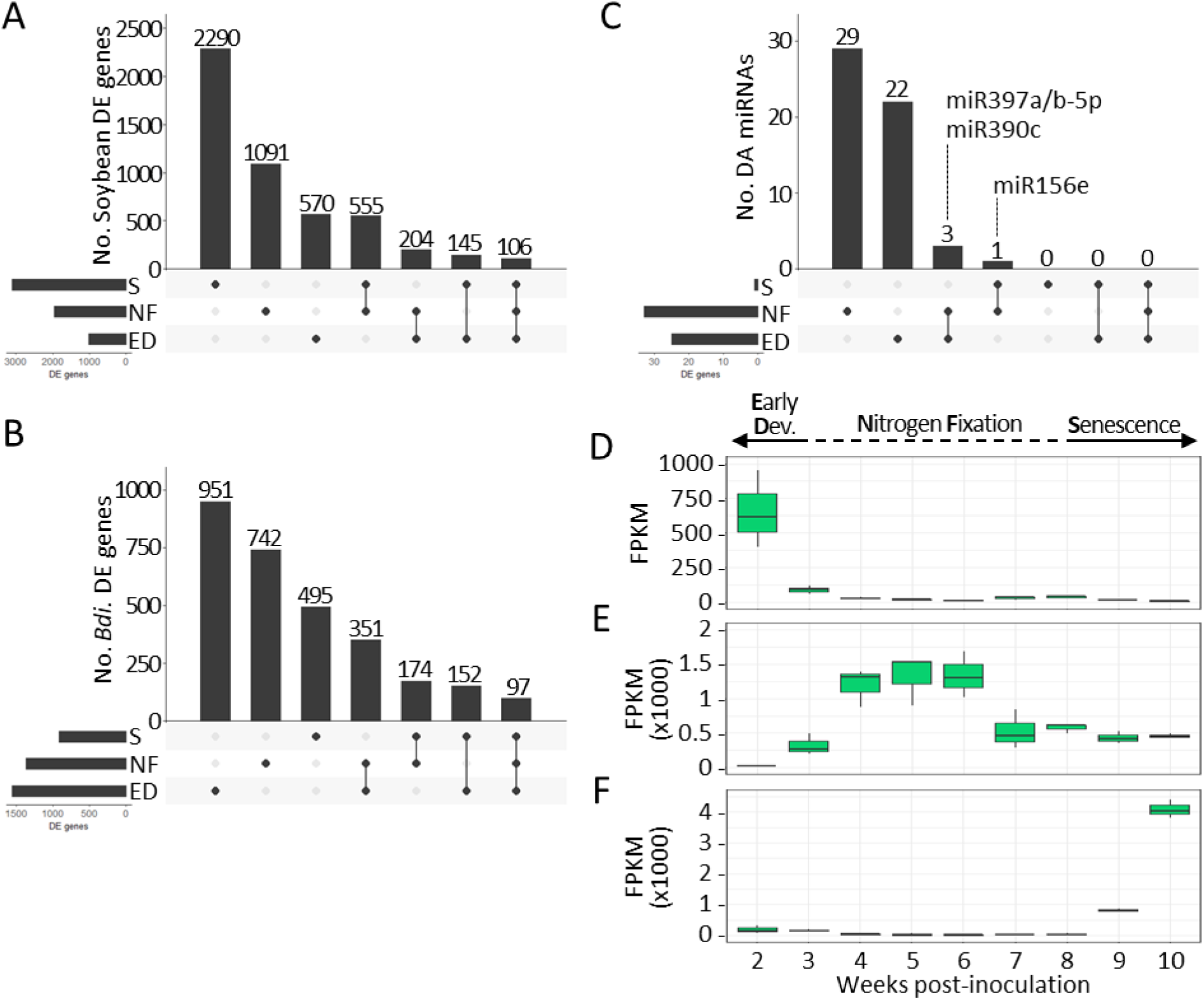
Stage-specific and multi-stage differential regulation of coding and non-coding transcripts during soybean nodule development (2 to 10 weeks post-inoculation) Upset plots illustrating stage-specific and multi-stage differential expression (DE) of genes and differential accumulation (DA) of miRNAs in soybean nodules. The bottom left panel shows the set sizes, the bottom right panel shows the intersection matrix, and the top right panel displays the number of DE genes or DA miRNAs in each combination. Stage-specific and shared DE genes in (A) soybean and (B) *Bradyrhizobium* across developmental stages (2 to 10 weeks post-inoculation): early development (ED, 2–3 wpi), nitrogen fixation (NF, 3–8 wpi), and senescence (S, 8–10 wpi). (C) Upset plot displaying unique and shared DA miRNAs across the same developmental stages. (D–F) Expression patterns of multi-stage DA miRNAs across the same developmental stages. Boxplots showing the expression of (D) miR390c, (E) miR397a/b-5p, and (F) miR156e, which exhibited multi-stage DA, but accumulation patterns suggest stage-specific roles in early nodule development, nitrogen fixation, and senescence, respectively.

Among stage-specific soybean DE genes, 58% (2,290/3,951) were present during senescence, reinforcing the extensive transcriptional reprogramming associated with nodule termination. The nitrogen fixation stage contained the next highest number of stage-specific DE genes, 28% (1,091/3,951), followed by early development with 14% (570/3,951). These findings suggest that soybean transcriptional activity increases progressively from early development (beginning at 2 wpi) through nitrogen fixation, and peaks during senescence (up to 10 wpi). However, as our study did not include time points prior to 2 wpi—a well-characterized period of symbiotic initiation and establishment—the apparent low number of DE genes during early development may reflect the tapering of earlier transcriptional activity, and it remains possible that early nodulation (0–2 wpi) exhibited DE gene numbers comparable to or even exceeding those observed during senescence.

In contrast, *Bradyrhizobium* exhibited its highest number of stage-specific DE genes during early development (43%, 951/2,188), followed by nitrogen fixation (34%, 742/2,188), and senescence (23%, 495/2,188). This pattern suggests that *Bradyrhizobium* undergoes its most extensive transcriptional reprogramming during early development, likely reflecting its adaptation to the nodule environment as it transitions from a free-living state to an endosymbiotic, nitrogen-fixing bacteroid. In contrast, soybean transcriptional regulation is most pronounced during senescence, coinciding with the host’s transition from flowering to pod-filling. This divergence underscores the host-driven nature of nodule termination, where soybean orchestrates the breakdown of symbiosis while *Bradyrhizobium* gradually transitions out of its symbiotic state.

During nitrogen fixation (3–8 wpi), soybean and *Bradyrhizobium* transcriptional changes stabilize (Figure 3), mirroring the metabolic equilibrium required for efficient nitrogenase activity. Despite spanning the longest developmental period (5-weeks), the nitrogen fixation stage contained fewer stage-specific DE genes per weekly comparison than either early development (one weekly comparison) in *Bradyrhizobium* (Figure 4B) or senescence (two weekly comparisons) in soybean (Figure 4A). This suggests that nitrogen fixation represents a transcriptionally stable phase, with minimal large-scale regulatory shifts compared to the sharp reprogramming events observed at the transitions into and out of symbiosis, early development and senescence, respectively.

For stage-specific DA miRNAs (Figure 4C), we identified 57% (29/51, 18 mature sequences) during nitrogen fixation, while the remaining 43% (22/51, 15 mature sequences) were DA during early development. Notably, we did not detect stage-specific DA miRNAs during senescence, suggesting that miRNA regulation primarily governs transitions into and out of the nitrogen fixation phase.

We also found that 22% of soybean DE genes (1,110/4,961) and 26% of *Bradyrhizobium* DE genes (774/2,962) were expressed in multiple stages. In soybean, half of these multi-stage DE genes (555/1,110) were active during both nitrogen fixation and senescence, highlighting major transcriptional changes during this transition. Another 18% (204/1,110) were expressed during early development and nitrogen fixation, marking the shift from symbiotic establishment to active nitrogen fixation. A smaller subset (13%, 145/1,110) was present during early development and senescence, suggesting a functional link between these stages, while the smallest group (10%, 106/1,110) spanned all three stages. Similarly, in *Bradyrhizobium*, 45% (351/774) of multi-stage DE genes were expressed during early development and nitrogen fixation. Smaller subsets (22% and 20%, 174/774 and 152/774, respectively) were expressed during nitrogen fixation and senescence or early development and senescence. A small fraction, 13% (97/774) was present in all three stages, suggesting broad regulatory roles. These multi-stage DE genes likely play key roles across different phases of nodule development and warrant further investigation.

In contrast, multi-stage DA miRNAs were rare; we identified just four of the 55 DA miRNAs of this type. Specifically, three miRNAs—miR390c, miR397/ab-5p, and miR156e—were present across two stages of development, with miR390c and miR397a/b-5p present during early development and nitrogen fixation, while miR156e was present during nitrogen fixation and senescence. However, closer examination of their accumulation patterns suggests distinct regulatory roles. miR390c was downregulated from 2 to 3 wpi and from 3 to 4 wpi **(Figure 4D)**. In contrast, miR397a/b-5p was upregulated from 2 to 3 wpi and from 3 to 4 wpi, reaching peak accumulation at 5 wpi before being downregulated from 6 to 7 wpi **(Figure 4E)**. These distinct patterns indicate that, despite both miRNAs being multi-stage DA during early development and nitrogen fixation, miR390c is enriched during early development, while miR397a/b-5p accumulated during nitrogen fixation. miR156e, DA during nitrogen fixation and senescence, showed low accumulation from 4 to 8 wpi before increasing sharply from 8 to 10 wpi, suggesting a role in senescence **(Figure 4F)**.

The identification of miR390c, miR397a/b-5p, and miR156e reinforce the three developmental stages identified in our transcriptomic analysis, with miR390c marking early development (2–3 wpi), miR397a/b-5p marking nitrogen fixation (3–8 wpi), and miR156e marking senescence (8–10 wpi). To further refine our understanding of nitrogen fixation and senescence regulation, we next investigated key genes and miRNAs involved in these developmental stages, identifying candidate molecular biomarkers as well as transcriptional regulators.

### Identification of seven novel candidate biomarkers for nitrogen fixation

To identify molecular markers for nitrogen fixation efficiency, we analyzed transcriptional patterns of soybean genes, plant miRNAs, and *Bradyrhizobium* genes associated with nitrogen fixation. Since nitrogenase activity peaked at 5 wpi, we focused on genes and miRNAs with peak expression at this stage. Weighted gene co-expression network analysis (WGCNA) identified 10 soybean gene modules (ME1-ME10) from a subset of 5,733 genes selected based on expression and variance thresholds. These modules represent genes with shared temporal expression trajectories across nodule development (2–10 wpi), highlighting distinct temporal co-expression patterns associated with symbiotic stages. **(Supplemental Table 11)**.

The two largest clusters, ME3 and ME2, accounted for ∼86% of identified genes **(Figure 5A)**. To classify developmental trends, we grouped soybean RNA expression modules by peak expression: ME3 (3,769 genes) and ME4 (68 genes) peaked 2–3 wpi (early development); ME1 (345 genes) peaked at both 5 and 8 wpi, suggesting a role post-peak nitrogen fixation; and ME2 (1,151 genes), ME7 (60 genes), and ME8 (73 genes) peaked at 8–10 wpi (senescence). Additionally, ME6 (124 genes) and ME10 (79 genes) exhibited biphasic expression from 4–7 wpi and 9–10 wpi, suggesting dual-roles in nitrogen fixation and senescence, while ME9 (20 genes) peaked at 2 and 6 wpi, indicating roles in distinct stages. ME5 (44 genes) had no clear pattern.

**Figure 5.**
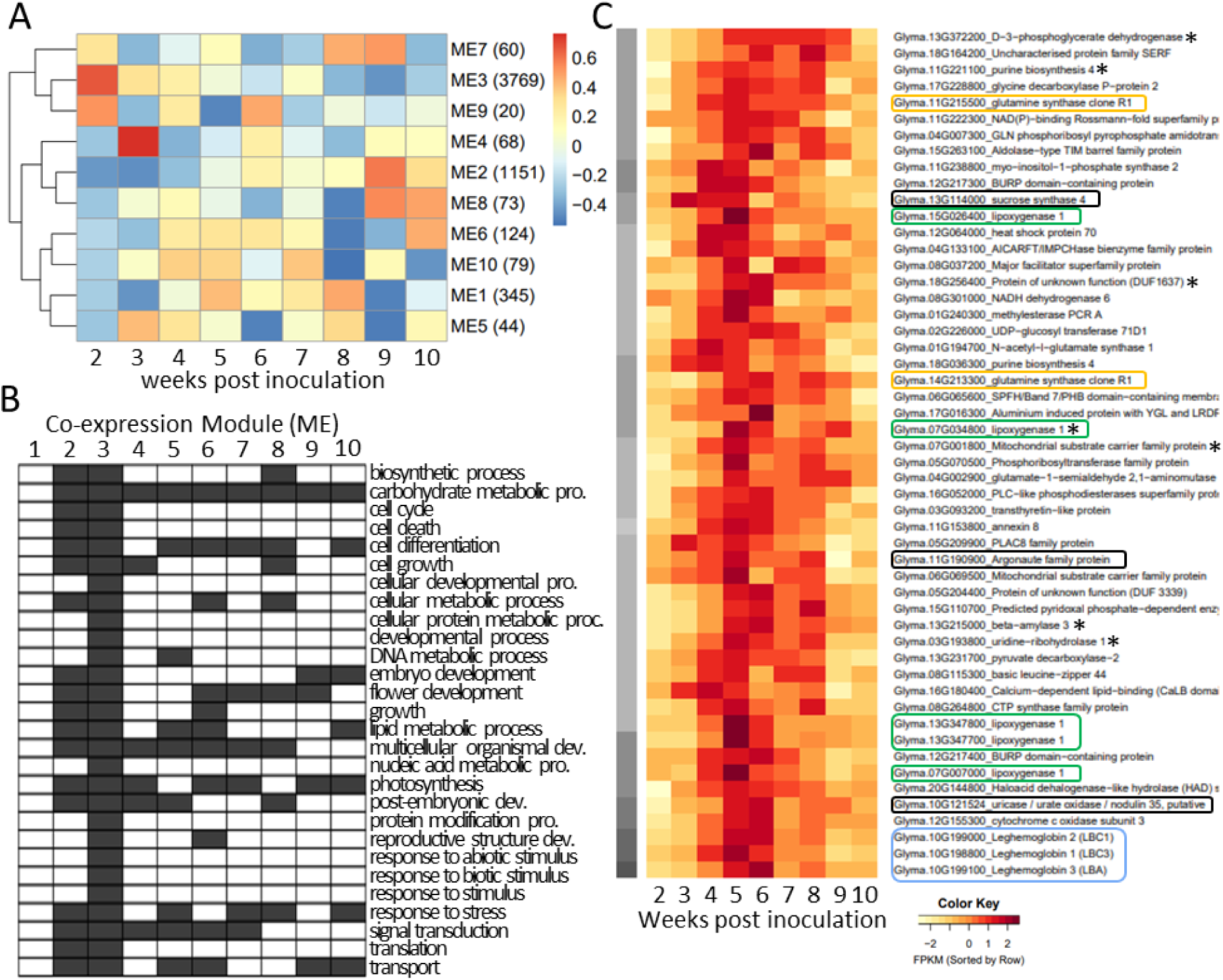
Identification of key soybean transcriptional biomarkers for nitrogen fixation in nodules. A) Weighted Gene Co-Expression Network Analysis (WGCNA) of soybean nodule transcriptomes. Hierarchical clustering of gene expression data grouped into 10 co-expression modules (ME1–ME10). The heatmap represents relative expression levels across nodule development (2–10 weeks post-inoculation, wpi), with high accumulation in red and low accumulation in blue. The dendrogram on the left indicates relationships between modules. (B) Gene Ontology (GO) analysis of WGCNA clusters using SoyBase (Grant et al. 2010). GO terms associated with biological processes are shown on the y-axis, with black boxes indicating presence within specific modules (x-axis). (C) Heatmap of highly expressed genes (FPKM > 1000) following a *2-up-5-down-10* pattern. Genes were upregulated (2–5 wpi) and downregulated (5–10 wpi), aligning with nitrogen fixation activity. Asterisks (*) mark seven genes significantly correlated with nitrogenase activity. Notable genes of interest include leghemoglobin (*Lb*; blue), *Lipoxygenase* (*Lox*; green), glutamine synthetase (*Gln*; yellow), and other key genes (black). Color intensity indicates expression level, with dark red representing high expression. Three biological replicates were analyzed per time point.

Gene Ontology (GO) analysis revealed key biological processes associated with WGCNA clusters **(Figure 5B)**. ME3, the largest module, contained genes spanning all GO categories, suggesting diverse biological activity during early development. Similarly, ME2, which peaked at 9 wpi (senescence), showed enrichment in multiple GO categories, suggesting that biological processes active in early development are reactivated or essential for nodule senescence. In contrast, nitrogen fixation appears to be governed by transient regulation, as no single ME peaked exclusively at 5 wpi. ME1, with minor peaks at 5 and 8 wpi, suggests that genes involved in nitrogen fixation are not confined to a single co-expression module but rather are distributed across multiple networks. Genes linked to carbohydrate metabolism were present in all modules but ME1, reinforcing their role throughout symbiosis. Additionally, stress response and stimulus perception genes were broadly distributed across senescence-associated modules (ME2, ME7, and ME8), highlighting complex regulatory changes as nodules shift from nitrogen fixation to termination.

Since WGCNA captured only ∼10% (5,733/56,044) of soybean genes and no single module peaked exclusively at 5 wpi, we performed targeted DE analysis to identify genes upregulated from 2 to 5 wpi and downregulated from 5 to 10 wpi. This “*2-up-5-down-10*” pattern aligns with nitrogen fixation activity, enabling biomarker identification. We identified 315 DE genes **(Supplemental Table 12)**, including 52 highly expressed genes (>1000 avg. FPKM from 2–10 wpi) **(Figure 5C)**. Among them, seven genes significantly correlated with nitrogenase activity (4–7 wpi) **(Supplemental Figure 5)**: *D-3-phosphoglycerate dehydrogenase* (Glyma.13G372200), *purine biosynthesis 4* (Glyma.11G221100), *lipoxygenase 1* (*Lox1*) (Glyma.07G034800), a protein of unknown function (DUF1637) (Glyma.18G256400), *uridine-ribohydrolase 1* (Glyma.03G193800), *beta-amylase 3* (Glyma.13G215000), and a *mitochondrial substrate carrier family protein* (Glyma.07G001800). These serve as strong candidate biomarkers for nitrogen fixation, warranting further characterization. Of the non-statistically significant *2-up-5-down-10* DE genes, *Leghemoglobins* (*Lb3, Lb2, Lb1*), ranked in descending order of expression, were among the top ten most highly expressed genes in nodules. These results reinforce their well-established role in oxygen buffering, essential for nitrogenase function and Bradyrhizobium respiration during symbiosis (Kuzma et al. 1993; Du et al. 2020).

### *Lipoxygenases* (*Lox*) genes as key nitrogen fixation biomarkers

We identified five *Lox* genes as highly expressed *2-up-5-down-10* DE genes, marking them as strong biomarkers for nitrogen fixation. Their role in nodules during nitrogen fixation remains unclear as they have not previously been identified in nodules. A study linking *Lox* to symbiosis described an inverse relationship between seed *Lox* levels and belowground nitrogen fixation (Zougari et al. 1995). *Lox* enzymes mediate oxylipin biosynthesis, which helps plants respond to stress (Singh et al. 2022). This may function to regulate oxygen balance in nodules by controlling lipid oxidation and ROS levels. Our results identified *Lox1* (Glyma.07G034800) as the only *Lox* gene significantly correlated with nitrogenase activity, while the other *Lox* genes did not meet the statistical threshold for correlation (Supplemental Figure 5). However, their shared expression pattern—peaking during nitrogen fixation and declining thereafter—suggests a functional role in symbiosis, potentially to maintain oxygen homeostasis in nodules during nitrogen fixation, warranting further investigation into their specific roles in SNF.

Several other notable genes exhibited the *2-up-5-down-10* expression pattern, highlighting their potential roles in nitrogen fixation. *Argonaute 5* (*AGO5*, Glyma.11G190900), a key regulator of small RNA pathways that primarily associates with 21- and 22-nucleotide sRNAs (Zhan and Meyers 2023), peaked at 5 wpi, reinforcing its previously established role in miRNA-mediated regulation of SNF (Reyero-Saavedra et al. 2017). *SUCROSE SYNTHASE 4* (*SUS4*, Glyma.13G114000), which cleaves sucrose into fructose and UDP-glucose, supplies the primary carbon source for bacteroid metabolism, ensuring rhizobial energy needs are met for efficient nitrogen fixation (Morell and Copeland 1985; Stein and Granot 2019). *SUS4* expression peaked early (3 wpi), remained stable through 5 wpi, and then declined, reflecting its essential role in fueling nitrogen fixation. Additionally, two glutamine synthetase genes, *Gln1;4* (Glyma.11G215500) and *Gln1;1* (Glyma.14G213300), displayed distinct expression patterns, highlighting their roles in SNF-derived nitrogen assimilation (Li et al. 2023). *Gln1;4* peaked at 5 wpi but remained highly expressed from 4 to 8 wpi, whereas *Gln1;1* peaked between 5 and 8 wpi before declining. Another gene, *Nodulin-35* (*N-35*, Glyma.10G121524), encoding a uricase involved in ureide metabolism (Nguyen et al. 2023), peaked at 6 wpi, suggesting a role in nitrogen redistribution as fixation declines. Although these genes did not significantly correlate with our study’s measurement of nitrogenase activity, their peak expression (4–6 wpi) suggests they support nitrogen fixation and nodule function.

### New miRNA biomarkers reveal a dynamic post-transcriptional regulation of SNF

To identify miRNAs associated with nitrogen fixation, we performed WGCNA and *2-up-5-down-10* DA analyses. WGCNA grouped 317 miRNAs into 11 modules (ME1-ME11) based on peak accumulation periods **(Figure 6A; Supplemental Table 13).** ME1 (120 miRNAs) and ME10 (20 miRNAs) peaked at 2 wpi (early development). ME7 (13 miRNAs) and ME8 (16 miRNAs) exhibited biphasic accumulation at 2 and 8 wpi or 2 and 7 wpi, suggesting roles in early development and senescence. ME2 (14 miRNAs) (5 wpi peak), ME3 (11 miRNAs) (3 and 7 wpi peaks), and ME6 (18 miRNAs) (6 wpi peak) were linked to nitrogen fixation. ME4 (38 miRNAs), ME5 (18 miRNAs), ME9 (16 miRNAs), and ME11 (14 miRNAs) peaked at 8–10 wpi (senescence). These clusters highlight dynamic miRNA regulation during nodule development. Among these, ME2 was the primary nitrogen fixation-associated module, peaking at 5 wpi, coinciding with peak nitrogenase activity. It contained 10 mature miRNA sequences from five canonical families—miR169 (miR169j-5p, miR169l-3p, miR169u, miR169v), miR391 (miR391-5p), miR397 (miR397a/b-5p, miR397b-3p), miR398 (miR398c/d), and miR399 (miR399a/b/c/h, miR399e/g), along with miR2111a/d. Filtering for highly accumulating miRNAs (>10 FPKM from 2–10 wpi) we identified six dominant miRNAs in descending order of expression: miR399a/b/c/h, miR397a/b-5p, miR169l-3p, miR169j-5p, miR398c/d, and miR2111a/d.

**Figure 6.**
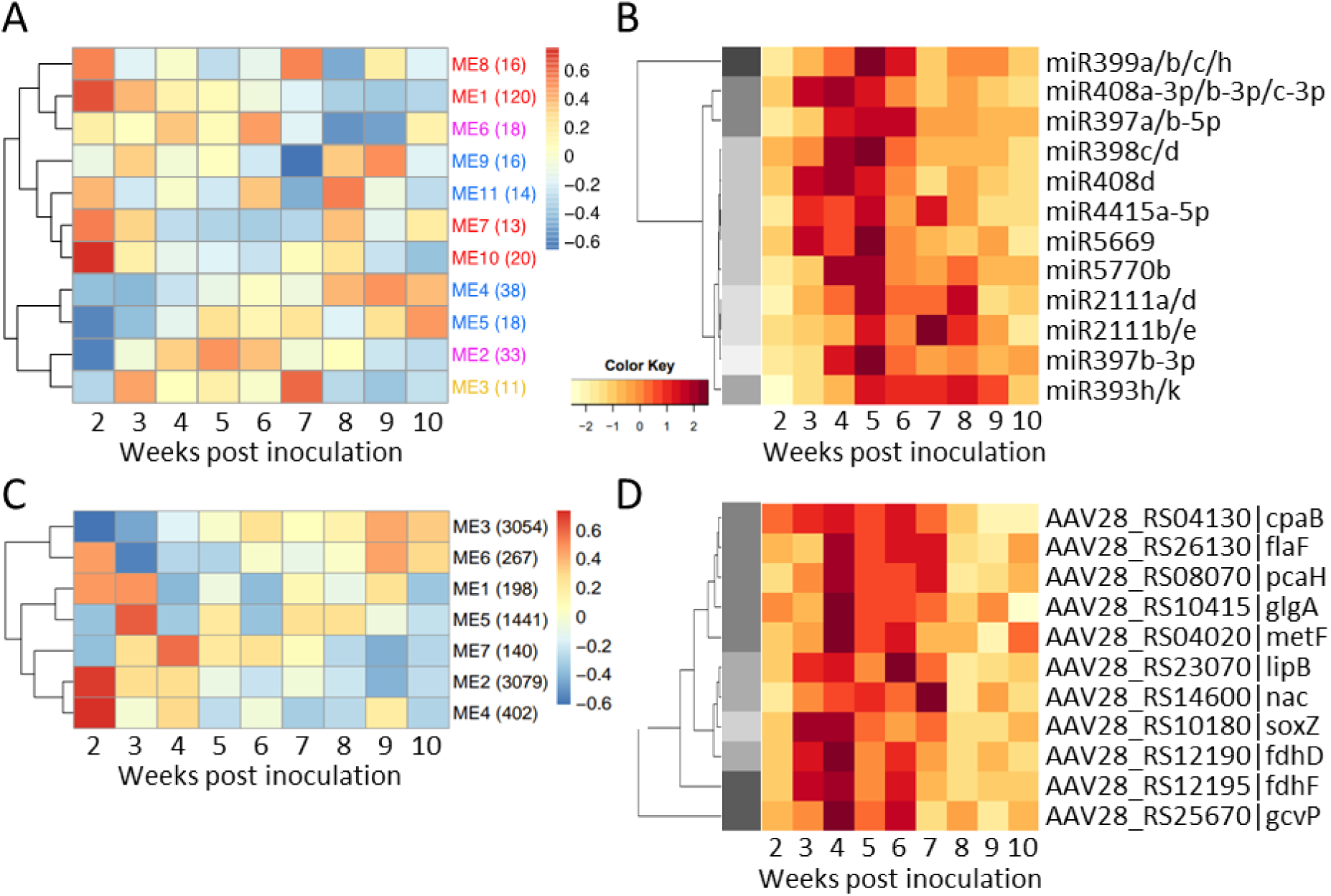
Identification of key miRNA and *Bradyrhizobium* biomarkers for nitrogen fixation using WGCNA and *2-up-5-down-10* differential expression patterns. (A) WGCNA clustering of soybean miRNAs across nodule development. Heatmap displaying 11 miRNA co-expression modules (ME1–ME11), grouped by peak accumulation. Colors indicate relative expression levels (high in red, low in blue). ME2, which peaks at 5 wpi, is most associated with nitrogen fixation. (B) Heatmap of highly expressed miRNAs following the *2-up-5-down-10* DA pattern. Color intensity represents relative expression, with dark red indicating peak accumulation. These miRNAs were upregulated from 2 to 5 wpi and downregulated from 5 to 10 wpi, aligning with nitrogenase activity. (C) WGCNA clustering of Bradyrhizobium genes across nodule development. Modules (ME1–ME7) are classified based on peak expression, with ME7 (4 wpi) identified as a key module for SNF-associated bacterial genes. (D) Heatmap of 11 annotated *Bradyrhizobium* genes identified in hierarchical clustering (WGCNA) ME7 exhibiting diverse expression patterns and peaking during nitrogen fixation.

The *2-up-5-down-10* DA analysis identified 22 miRNAs following the nitrogen fixation timeline, including members of five major families—miR393, miR397, miR398, miR399, and miR408—as well as miR2111 **(Figure 6B; Supplemental Table 14).** These families regulate nutrient signaling, oxidative stress, plant growth, and rhizobial infection, all of which are critical for efficient nitrogen fixation (Cai et al. 2017; De Luis et al. 2012; Li et al. 2022a; Fan et al. 2021; Zhang et al. 2021). Notably, all but miR393 and miR408 were also present in ME2, reinforcing their relevance in SNF. Previously characterized in shoots and roots, with a role in AON (Tsikou et al. 2018; Okuma and Kawaguchi 2021), we observed miR2111a/d and miR2111b/e in nodules, peaking at 5 wpi and 7 wpi, respectively. These results suggest potential new roles for miR2111 in coordinating nitrogen fixation, integrating systemic and local symbiotic regulation within nodules. Other notable families include miR393, which regulates auxin signaling by targeting TIR1/AFB auxin receptors and modulating Aux/IAA degradation (Jiang et al. 2022), accumulated from 5–8 wpi, suggesting a role during nitrogen fixation, deviating from its known role in roots during early development, in which it regulates nodulation via its target, *ENOD93* (Yan et al. 2015; Cai et al. 2017). miR397, which targets laccase genes involved in lignin biosynthesis and nodule structure (Huang et al. 2021), peaked from 4–6 wpi, suggesting a role in maintaining nodule structural integrity during periods of elevated nitrogen fixation activity. These findings reinforce miR397 as a regulator of nodule function and nitrogen fixation in determinate-nodule forming legumes, such as soybean and *Lotus japonicus* (De Luis et al. 2012). Additionally, miR398, previously identified to be upregulated in mature nodules (Yan et al. 2015), was highly enriched in 5 wpi nodules in our study. A known regulator of oxidative stress, miR398 is known to target *Cu/Zn superoxide dismutase* (CSD) genes, a highly conserved gene family in plants, responsible for scavenging reactive oxygen species (ROS) and mitigating oxidative stress (Li et al. 2022a). The enrichment of miR398 suggests a localized decline in nodule ROS inhibition during or immediately following peak nitrogen fixation, with prior and subsequent periods of higher activity (Figure 6B). We also detected miR399, the most highly accumulated miRNA exhibiting the *2-up-5-down-10* pattern, and miR408, both key regulators of phosphate signaling and ATP metabolism (Xu et al. 2013). Their presence in nodules at 5 wpi suggests a potential role in linking phosphate homeostasis to SNF energy balance. Notably, miR399, has been previously linked to nitrogen fixation efficiency, with its induction proposed to enhance both nitrogen fixation and soybean growth (Fan et al. 2021). This hypothesis is further supported by our observations, where miR399a/b/c/h was enriched in nodules during peak nitrogen fixation, reinforcing its potential role in optimizing SNF.

Together, these findings highlight miRNA-based regulatory networks governing nitrogen fixation. The combined WGCNA and DA analyses identified multiple miRNAs peaking during nitrogen fixation, suggesting key roles in nodule function and symbiotic efficacy. The distinct accumulation patterns and levels emphasize the complexity of miRNA regulation, and suggest cell-type-specific functions as possible explanations. Future studies using single-cell transcriptomics could resolve miRNA accumulation patterns across nodule cell types, providing deeper insight into miRNA-mediated SNF regulation.

### *Bradyrhizobium cpaB* and *gcvP* genes play critical roles in nitrogen fixation and symbiotic efficiency

To identify *Bradyrhizobium* genes linked to SNF, we did the same combined differential gene analysis. The *2-up-5-down-10* DE analysis yielded 12 genes, none of which were annotated **(Supplemental Table 15)**. We observed that *Bradyrhizobium* undergoes the most transcriptional changes during pre-peak nitrogen fixation (2–5 wpi) and senescence (8–10 wpi), suggesting that host developmental transitions, rather than nitrogen fixation activity, dictate bacterial gene regulation. WGCNA clustered *Bradyrhizobium* transcripts into seven co-expression modules (ME1–ME7) **(Figure 6C; Supplemental Table 16)**. We classified modules by peak expression: ME1 (198 genes), ME2 (3,079 genes), ME4 (402 genes), ME5 (1,441 genes), and ME7 (140 genes) peaked at 2–4 wpi, aligning with early development. ME6 (267 genes, peaking at 2 and 9 wpi) exhibited biphasic expression, suggesting roles in early nodule adaptation and senescence. ME3 (3,054 genes, peaking at 9 wpi) was linked to nodule senescence. ME2 and ME3, accounting for ∼71.5% of clustered transcripts, primarily represented genes involved in early development and senescence, respectively. These results align with broader DE trends showing minimal bacterial transcriptional enrichment during nitrogen fixation (5–8 wpi).

No *Bradyrhizobium* module peaked at 5 wpi, but ME7 peaked at 4 wpi. This, along with a small number of genes identified by *2-up-5-down-10* DE analysis, suggests that bacterial genes essential for nitrogen fixation are activated prior to peak nitrogenase activity, making ME7 a key module for SNF biomarker identification. Among ME7 genes, 11 were annotated, three of which—*lipB*, *nac*, and *soxZ*—exhibited stable or minimally fluctuating expression across nodule development, suggesting they may play constitutive roles throughout symbiosis rather than being specifically involved in SNF **(Figure 6D)**.

Among the other eight, *cpaB*, is part of the colonization pilus (*cpa*) operon and facilitates bacterial attachment and biofilm formation (Streit et al. 2004). Its expression pattern (Supplemental Table 3) suggests that adherence within nodule cells is crucial for SNF efficiency. Additionally, its gradual decline in expression from 6–10 wpi may be triggered by reduced structural attachment needs as nodules transition toward senescence and bacteria shift towards a post-symbiotic, free-living state. Similarly, *flaF,* which encodes a flagellar basal-body component, is involved in bacterial surface structure (Sandhu et al. 2023). However, despite showing peaks on the heatmap, *flaF* expression was highly variable among biological replicates, exhibiting a fluctuating pattern without a clear peak.

The remaining six genes—*gcvP, fdhF, metF, glgA, pcaH,* and *fdhD* (listed in descending order of average expression (2–10 wpi)—peaked at 4 wpi. Among them, *gcvP* (*glycine decarboxylase*) had the highest overall average expression, 2.1 times higher than *fdhF*. The system of proteins encoded by *glycine cleavage* (*gcv*) genes play a central role in one-carbon metabolism and energy transfer (Kikuchi et al. 2008) and influence soybean cultivar specificity (Lorio et al. 2010). Inactivating *gcvP* in *Sinorhizobium fredii* USDA257 enabled it to nodulate North American soybean cultivars previously incompatible with infection, suggesting a role in host compatibility and nodulation efficiency. The presence of *fdhF* and *fdhD* (formate dehydrogenases) highlights their role in electron transport during nitrogen fixation, supplying reducing power in the form of NADH for nitrogenase function in the microaerobic nodule environment (Wang and Gunsalus 2003). Similarly, *metF* (methylenetetrahydrofolate reductase) contributes to methionine biosynthesis and one-carbon metabolism, essential for nitrogen assimilation and bacterial metabolic regulation during SNF (Price et al. 2021). The identification of *glgA* (glycogen synthase) suggests glycogen accumulation as a carbon storage strategy in bacteroids. This may sustain nitrogen fixation under fluctuating carbon availability, ensuring metabolic flexibility during symbiosis (Wang et al. 2007). Additionally, *pcaH* (protocatechuate 3,4-dioxygenase subunit) plays a role in aromatic compound metabolism, mitigating oxidative stress, and maintaining redox balance (Ohlendorf et al. 1988)—likely critical for nitrogenase function.

### Identification of six novel soybean senescence biomarkers (*GS-ASN1, Hb2, Lea3-5, NF-YA3, PDS1*, and *SUS3*) in nodules

Senescence marks the final stage of nodule development, terminating SNF and reallocating resources to the plant. This process involves significant morphological, physiological, and biochemical changes, yet the molecular mechanisms driving nodule senescence remain largely underexplored (Dupont et al. 2012; Kazmierczak et al. 2020; Zhou et al. 2021). To pinpoint soybean senescence-associated genes, we examined WGCNA clusters enriched in late-stage nodules (8–10 wpi) (Figure 5A). Three co-expression modules—ME2, ME7, and ME8—peaked between 8 and 10 wpi. While ME7 and ME8 also contained genes active earlier, ME2 was predominantly enriched during senescence, making it the most relevant for identifying late-stage biomarkers.

### Identification of potential miR2111-*TML* regulatory interactions in nodules

Within ME2, we identified *RIC1* (Glyma.13G292300) and *TML2* (Glyma.05G077700), previously characterized as AON regulators in roots during early stages of nodulation (Reid et al. 2011; Wang et al. 2019; Tsikou et al. 2018; Chaulagain et al. 2023; Gresshoff et al. 2023), suggesting potential roles in local nodule termination (Supplemental Table 2). Additionally, the enrichment of miR2111 family members—known to target soybean *TML* genes in roots (Zhang et al. 2021)—during nitrogen fixation (Figure 6B) prompted us to examine the expression patterns of two other soybean *TML* genes, *TML1a* (Glyma.16G057300) and *TML1b* (Glyma.19G090600). All three *TML* genes exhibited a high-to-low-to-high expression pattern, with peaks at 2 and 10 wpi, indicative of potential roles in both early development and senescence. While *TML2* displayed relatively low expression, its most pronounced peak occurred at 10 wpi, whereas *TML1a* and *TML1b* showed higher transcript levels, with *TML1a* being the most highly expressed, peaking most prominently at 2 wpi (Supplemental Table 2). These expression patterns suggest potential miR2111-*TML* regulatory interactions within nodules and warrant further investigation.

To refine our candidate list, we conducted a “*2-up-10*” DE analysis, reducing the pool from 1,151 to 453 genes that exhibited sustained upregulation during senescence **(Supplemental Table 17)**. Further filtering for high expression (>600 avg. FPKM, 8–10 wpi) yielded 21 genes, nearly all peaking at 9 wpi before declining at 10 wpi **(Figure 7A)**. Notably, 12 of these displayed a biphasic expression pattern, with a minor peak at 6 wpi, dropping slightly at 7 wpi, and rising again at 9 wpi before declining at 10 wpi, suggesting roles in nodule aging and senescence. We observed this trend in *Lea3-5*, *SUS3*, *NF-YA3*, *Hb2*, and *PDS1*, among others. *Late embryogenesis abundant-3_5 (Lea3_5*, Glyma.10G017600), encoding a member of the LEA protein family known for stress tolerance, is typically linked to desiccation and osmotic protection (Shiraku et al. 2022; Guo et al. 2023). Its upregulation during senescence suggests a protective role as nodules mature and undergo structural and metabolic shifts associated with aging. *Hemoglobin 2* (*Hb2*, Glyma.11G121800), traditionally classified as non-symbiotic and believed to originate from non-infected cells within nodules (Anderson et al. 1996; Hill et al. 2016; Du et al. 2020), exhibited increasing expression from 2 to 9 wpi, suggesting involvement in oxidative stress regulation or metabolic adaptation as SNF declines. *PHYTOENE DESATURASE-1* (*PDS1,* Glyma.14G030400), involved in carotenoid biosynthesis, was significantly upregulated at 9 wpi. Carotenoids protect cells from oxidative damage, positioning *PDS1* as a potential player in antioxidative defense during nodule aging, a largely unexplored function in nodules.

**Figure 7.**
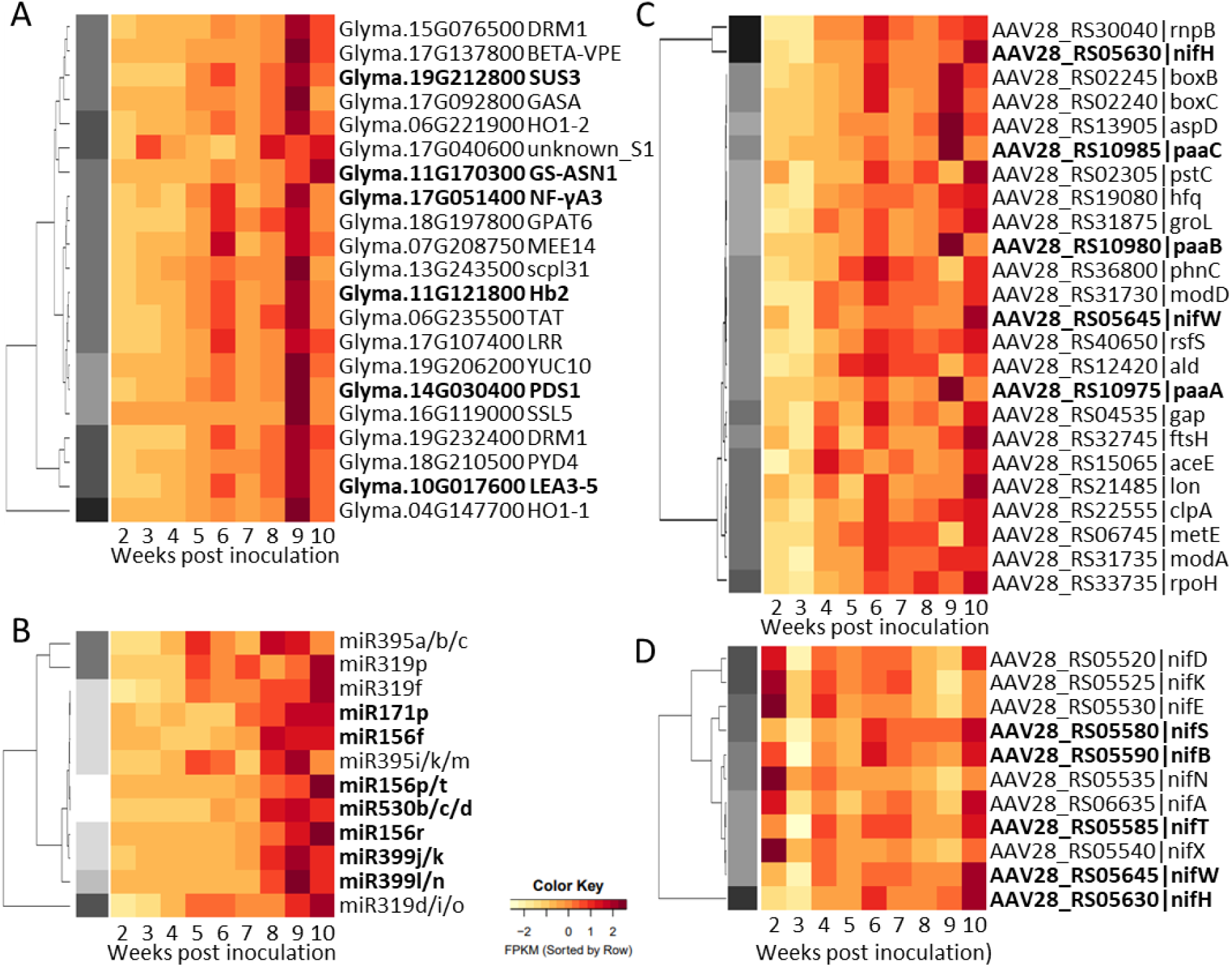
Identification of soybean and *Bradyrhizobium* senescence biomarkers through co-expression and *2-up-10* differential expression analysis. (A) Heatmap of 21 highly expressed soybean senescence-associated genes identified by *2-up-10* differential expression analysis. These genes were upregulated from 2 to 10 wpi, with most peaking at 9 wpi before declining. Several genes, including *Lea3-5, SUS3, NF-YA3, Hb2,* and *PDS1*, displayed biphasic expression, suggesting roles in oxidative stress regulation, carbohydrate remobilization, and nitrogen remobilization during nodule aging. *GS-ASN1* exhibited a continuous rise in expression from 4 to 10 wpi, highlighting it as a strong senescence biomarker. (B) Heatmap of 12 senescence-associated miRNAs identified by 2-up-10 differential accumulation analysis. These miRNAs exhibited sustained accumulation from 8 to 10 wpi, with some showing a distinct low-to-high accumulation pattern across nodule development (miR156f, miR156p/t, miR156r, miR171p, miR399j/k, miR399l/n, and miR530b/c/d), indicating stage-specific regulatory roles. (C) Heatmap of 24 highly expressed *Bradyrhizobium* genes upregulated from 2 to 10 wpi, including *nifH* and *nifW*. These genes exhibited peak expression at 10 wpi, suggesting continued bacterial metabolic activity and adaptation as host resources decline. (D) Expression dynamics of *Bradyrhizobium nif* genes across nodule development. *nifK, nifE, nifN, nifX* peaked exclusively during early development, while *nifH, nifW, nifT,* and *nifS* were exclusively upregulated at 10 wpi, indicating roles in nodule aging and senescence. Alternatively, *nifD* and *nifA* exhibited peaks during early development (2 wpi) and senescence (10 wpi), while *nifB* showed three peaks at 2, 6, and 10 wpi.

We also identified *SUS3,* (Glyma.19G212800), suggesting divergent *SUS* gene functions in nodules. While *SUS4* (Glyma.17G171700) was enriched during nitrogen fixation, likely fueling bacteroid metabolism, *SUS3* may facilitate host carbohydrate remobilization during senescence (Morell and Copeland 1985; Gordon et al. 1999; Stein and Granot 2019). *NF-YA3* (Glyma.17G051400), a transcriptional regulator of *Early Nodulation* (*ENOD40*), targeted by miR169c (Xu et al. 2021), was upregulated in senescence (9 wpi peak) as miR169j-5p and miR169l-3p were rapidly downregulated at 6 wpi and 7 wpi, respectively (Supplemental Table 4). Given its role in nitrogen-responsive transcriptional regulation (Leyva-González et al. 2012), increased *NF-YA3* expression may contribute to nitrogen remobilization processes during nodule decline.

Unlike other senescence-associated genes that peaked at 9 wpi before declining, *GS-ASN1* (Glyma.11G170300), encoding a glutamine-hydrolyzing asparagine synthetase, exhibited a unique and continuous increase in expression from 2 to 10 wpi, reaching its highest levels at 10 wpi (Figure 7A). This expression pattern distinguishes it as the only highly expressed *2-up-10* DE gene that did not decline at 10 wpi, underscoring its potential importance in late-stage nodule function. *GS-ASN1* has not been previously identified in nodules or symbiosis-related studies but was recently found to be differentially expressed in soybean leaves in response to environmental conditions (Hooker et al. 2023). Given the critical role of asparagine synthetases in nitrogen remobilization, its sustained upregulation suggests a key function in redistributing nitrogen as SNF ceases. Its steady accumulation from 4 wpi onward, with a continuous upward trajectory, positions *GS-ASN1* as a more robust senescence biomarker than previously identified genes.

These findings establish *GS-ASN1, Hb2, Lea3-5, PDS1, NF-YA3,* and *SUS3* as key senescence biomarkers, implicate asparagine and carotenoid biosynthesis in nodule aging, and introduce a potential miR2111*-TML* regulatory mechanism in nodules. To further elucidate the functions of these genes within symbiosis, single-cell transcriptomic technologies, such as single-nucleus RNA-seq (Cervantes-Pérez et al. 2024), could aid in resolving the cellular origins of these genes (i.e. infected vs. non-infected cell types) within nodules.

### Senescence-Associated miRNAs Fine-Tune Transcriptional Programs and Sustain Nodule Aging

In previous analyses, we identified 86 miRNAs from four WGCNA groups (ME4, ME5, ME9, and ME11) that had high accumulation from 8 to 10 wpi, suggesting roles in nodule aging and senescence (Figure 6A). Of these, 34 were DA and upregulated from 2 to 10 wpi, 23 of which belonged to six canonical families: miR156, miR171, miR319, miR395, miR399, and miR530 **(Figure 7B; Supplemental Table 18)**. This set included 12 mature miRNA sequences: miR156f, miR156p/t, miR156r, miR171p, miR319d/i/o, miR319f, miR319p, miR395a/b/c, miR395i/k/m, miR399j/k, miR399l/n, and miR530b/c/d. Unlike coding genes, which exhibited definitive peaks at 9 wpi (Figure 7A), senescence-associated miRNAs showed a more sustained accumulation from 8 to 10 wpi, suggesting a prolonged regulatory role. Seven of these: miR156f, miR156p/t, miR156r, miR171p, miR399j/k, and miR399l/n showed a distinct low to high accumulation pattern (2–10 wpi), with prominent accumulation from 8 to 10 wpi, indicating stage-specific roles in nodule aging and senescence. In contrast, the other five miRNAs―miR395a/b/c, miR319p, miR319f, miR395i/k/m, and miR319d/i/o―exhibited biphasic accumulation patterns suggesting a minor role in nitrogen fixation (peaking at 5 wpi), followed by a more prominent role in senescence (8–10 wpi peaks). We also observed distinct miRNAs from the same families with accumulation patterns in different stages of development (Supplemental Table 13). For example, miR171p accumulates during senescence, while 10 miR171 members regulate early development (ME1 and ME10), and two show biphasic accumulation during early development and senescence (ME7). Similarly, the miR399 family contains members enriched exclusively during either nitrogen fixation, such as miR399a/b/c/h (Figure 6B) or during senescence, as we observed with miR399j/k and miR399l/n (Figure 7B). The miRNAs miR156p/t and miR530b/c/d accumulated gradually from 2 to 10 wpi but remained low (<10 avg. FPKM at 8–10 wpi), suggesting a minor or cell-specific role within senescing nodules. Taken together, our miRNA analysis indicates that some miRNA families, such as miR171 and miR399, contain individual members that play specific roles during discrete stages of nodule development.

### *Bradyrhizobium* transcriptional reprogramming during nodule senescence signals a shift toward free-living survival

As nodules approach senescence, *Bradyrhizobium* undergoes transcriptional shifts, transitioning from a nitrogen-fixing bacteroid to a survival-oriented free-living state. We identified two senescence-associated bacterial co-expression modules: ME3 and ME6 (Figure 6C). ME3 gradually increased (5–9 wpi), peaking at 10 wpi with a minor peak at 6 wpi, mirroring senescence-associated soybean genes. ME6 exhibited biphasic expression, peaking at 2 and 9 wpi, suggesting roles in early development and senescence. Since this analysis focused on bacterial responses to senescence, we examined stage-specific genes within ME3. Of its 3,054 genes, 1,936 were DE and upregulated from 2 to 10 wpi, with 154 annotated. From those, 24 genes were highly expressed from 8 to 10 wpi (>1,000 avg. FPKM), including *nifH* and *nifW* **(Figure 7C; Supplemental Table 19)**.

Since *nif* genes have previously been described in nitrogen fixation (Frank et al 2018), we examined *nif* gene expression in our samples. Our analysis revealed two distinct groups: early development and senescence **(Figure 7D)**. Early-expressed *nifA, nifD, nifE, nifK, nifN*, and *nifX* (ME2) peaked at 2 wpi before declining but remained stable across development. The genes *nifD, nifA*, and *nifX* had secondary peaks at 10 wpi, while *nifK* and *nifN* showed minor peaks at 4 wpi. In contrast, *nifH, nifW, nifB, nifS,* and *nifT* (ME3) peaked at 10 wpi, with a minor peak at 6 wpi, suggesting roles in both nitrogen fixation and senescence. Among *nif* genes, *nifH* was the highest-expressing and the third most abundant *Bradyrhizobium* expressed gene in nodules (2–10 wpi), consistent with previous studies (Franck et al. 2018). Rhizobial *nifH* encodes the Fe-protein subunit of nitrogenase, transferring electrons to the MoFe-protein complex (Hu and Ribbe 2013). Additionally, *nifD* and *nifK*, encoding MoFe subunits, were the fourth and fifth most highly expressed genes, reinforcing their essential SNF roles. The sustained expression of *nifH, nifD*, and *nifK* highlights their role in nitrogenase activity while also suggesting continued involvement in senescence as *Bradyrhizobium* transitions to a free-living state. While *nifD* and *nifK* peak earlier, the gradual increase in *nifH* suggests distinct regulatory mechanisms, potentially extending its function beyond active nitrogen fixation.

Beyond *nif* genes, several highly expressed ME3 genes suggest metabolic adaptation and stress responses during senescence. Three genes encoding phenylacetic acid (PAA) degradation components—*paaA, paaB,* and *paaC*—were highly enriched in senescing nodules. The *paa* operon, which encodes enzymes that degrade phenylacetic acid (Cafiero et al. 2023), likely provides rhizobia with an alternative carbon source under nutrient-limited conditions, such as those encountered during senescence. Its upregulation at 9 wpi implies that *Bradyrhizobium* shifts to alternative carbon sources as host resources decline during senescence. Additionally, *rpoH*, encoding a heat shock sigma factor, was among the most highly expressed genes in ME3 (Figure 6C). *RpoH* regulates responses to heat and oxidative stress, controlling chaperones and proteases that mitigate protein misfolding (Martínez-Salazar et al. 2009). Its sustained expression from 2 to 10 wpi suggests an active bacterial response to proteotoxic stress as the nodule microenvironment ages.

These findings indicate that *Bradyrhizobium* remains transcriptionally and metabolically active during senescence, balancing a decline in support from the host with the need for survival beyond symbiosis. The sustained expression of *nifD, nifK,* and *nifH*, along with genes involved in alternative carbon metabolism (*paa* operon) and stress adaptation (*rpoH*), underscores a dynamic bacterial response, where *Bradyrhizobium* persists even as nitrogen fixation declines, suggesting a transition to a free-living state post-symbiosis.

## DISCUSSION

The regulation of nitrogen fixation in soybean nodules involves complex transcriptional and post-transcriptional controls that orchestrate symbiotic establishment, maintenance, and senescence. Our transcriptomic analyses revealed that while soybean undergoes extensive transcriptional reprogramming during early development and senescence, *Bradyrhizobium* exhibits its most significant regulatory shifts early in symbiosis, reinforcing the host-driven nature of nodule development. The identification of three major developmental phases—early development (2–3 wpi), nitrogen fixation (3–8 wpi), and senescence (8–10 wpi)—underscores the dynamic and tightly controlled temporal nature of this symbiotic interaction.

Among key regulatory elements, the identification of miR2111 family members and their potential *TML* targets in nodules indicates a putative regulatory pathway linking systemic AON with local nitrogen fixation control. Six miR2111 family members, represented by two mature sequences, miR2111a/d and miR2111b/c/e/f, exhibited peak accumulation at 5 wpi, coinciding with maximum nitrogenase activity. In systemic AON, shoot-derived miR2111 suppresses root-expressed *TML* genes to regulate nodule number (Zhang et al. 2021). Our identification of *TML1a, TML1b,* and *TML2* transcripts in nodules, all displaying high-to-low-to-high expression patterns (inverse to miR2111 accumulation) suggests that miR2111 may also function locally within nodules, potentially regulating *TML* activity during nitrogen fixation. However, despite the detection of miR2111 and *TML* transcripts in nodules, their precise origin remains unclear, as miR2111 is known to be transported systemically from shoots (Tsikou et al. 2018). Further research is required to determine whether these transcripts and miRNAs are locally expressed in infected or uninfected nodule cell-types or are remnants of systemic signaling. Future studies should focus on single-cell spatial expression profiling and functional validation to clarify the extent to which miR2111-*TML* interactions regulate nodule development and nitrogen fixation within nodules.

miR169 family members exhibited differential accumulation patterns, suggesting functional divergence in their roles during symbiosis. In particular, miR169j-5p and miR169l-3p reached peak accumulation from 5–6 wpi, before rapidly declining at 7 wpi, coinciding with the onset of nodule senescence and an expected decline in nitrogen fixation. Notably, *NF-YA* transcripts, known targets of miR169 in soybean (Xu et al. 2021; Ni et al. 2013), exhibited inverse expression patterns, remaining low during peak nitrogen fixation before steadily increasing following declines in miR169j-5p and miR169l-3p. The *NF-YA* transcription factor family has been implicated in nitrogen-responsive transcriptional regulation in Arabidopsis and maize (Leyva-González et al. 2012; Luan et al. 2014) and plays a critical role in regulating nodulation in legume-*Rhizobium* symbiosis (Zanetti et al. 2017). Here, we observed *NF-YA3* (Glyma.17G051400) with a low-to-high expression pattern, suggesting an additional role in nodule senescence. The reciprocal expression dynamics of miR169 and *NF-YA3* indicate that specific miR169 members may act as developmental regulators, integrating nitrogen availability cues to fine-tune nodule *NF-YA* expression at key transition points rather than directly regulating nitrogen fixation. This supports the idea that nitrogen fixation is not controlled by a single regulatory switch but is finely coordinated between host developmental cues and environmental nitrogen status (Lepetit and Brouquisse 2023).

The identification of *Lox1* (*Glyma.07G034800*) and other lipoxygenase (*LOX*) genes as potential biomarkers for nitrogen fixation suggests a novel role for oxylipin metabolism in nodules. Lipoxygenases are iron-containing dioxygenases that catalyze the oxygenation of polyunsaturated fatty acids, generating signaling molecules involved in plant development and stress responses (Andreou and Feussner 2009). While their roles in symbiosis remain largely unexplored, their upregulation during nitrogen fixation suggests functions beyond primary metabolism. One possibility is that *LOX* enzymes contribute to oxygen homeostasis within nodules, facilitating nitrogenase activity by assisting leghemoglobin in maintaining an optimal microaerobic environment. Alternatively, oxylipins produced by *LOX* genes may regulate host responses to nitrogen fixation, influencing nodule longevity and their disappearance, marking the transition to senescence. Given the close correlation between *Lox1* expression and nitrogenase activity, further studies should investigate whether oxylipin signaling is integral to SNF regulation.

*Bradyrhizobium* transcriptional shifts during senescence indicate a metabolic transition toward survival beyond symbiosis. The PAA operon gene family (*PAA*), which encodes enzymes for phenylacetic acid degradation, was significantly upregulated at 9–10 wpi. *PAA* catabolism provides an alternative carbon source under nutrient-limited conditions and is associated with bacterial stress responses (Cafiero et al. 2023). Its increased expression in senescing nodules suggests that *Bradyrhizobium* adapts its metabolism to declining host carbon supply, supporting its persistence in deteriorating symbiotic conditions. Similarly, the upregulation of *rpoH*, encoding a heat shock sigma factor, highlights bacterial responses to proteotoxic stress in senescing nodules. *RpoH* regulates genes involved in protein folding, repair, and degradation, ensuring rhizobial cellular integrity under stress conditions (Martínez-Salazar et al. 2009). The sustained expression of *nifH* and other key *NIF* genes alongside these survival-associated genes further suggests that nitrogenase components may serve additional functions beyond SNF, potentially aiding in bacterial adaptation to nodule senescence.

The observed period of transcriptional homeostasis from 5 to 6 wpi, marked by minimal differentially expressed genes and miRNAs in both soybean and *Bradyrhizobium*, raises intriguing questions about the regulatory mechanisms governing nitrogen fixation maintenance. The detected shift from 22-nt sRNAs enriched in early developing nodules (2 wpi) to 21-nt sRNAs from 4 wpi onwards, suggests a developmental transition in sRNA-mediated regulation, potentially linked to the onset of nitrogen fixation and modulation of symbiotic gene expression. Additionally, despite stability in soybean and *Bradyrhizobium* DE transcript levels (5–8 wpi), miRNA differential accumulation increased at 6–7 wpi, suggesting that post-transcriptional mechanisms may contribute to nitrogen fixation regulation during this phase. Furthermore, the delayed response of host and bacterial gene expression to miRNA activity implies that miRNA-mediated control may primarily function through translational repression rather than immediate transcript degradation (Iwakawa and Tomari 2013). This phase of transcriptional stability could signify a metabolic equilibrium, where symbiosis operates at peak efficiency with minimal regulatory adjustments before declining nitrogenase activity during senescence.

Our findings also provide novel insights into the role of soybean hemoglobins in nodules. We detected *Hb2*, previously classified as a non-symbiotic hemoglobin, in senescing nodules. Unlike classical leghemoglobin, which facilitates oxygen transport to support nitrogenase activity (Du et al. 2020), *Hb2* may be involved in oxidative stress mitigation during nodule aging. As leghemoglobin expression was low in senescing nodules (8–10 wpi) (Figure 5C), its lowered quantity likely alters the oxidative environment, increasing ROS accumulation and triggering *Hb2* expression. Hemoglobins have been shown to respond to abiotic stress by scavenging reactive oxygen and nitrogen species (ROS and RNS), which accumulate in senescing nodules (Gupta et al. 2011; Koltun et al. 2022; Puppo et al. 2005), potentially protecting both plant and bacterial cells from damage. The discovery of *Hb2* in nodules indicates that its function extends beyond non-symbiotic roles, opening new avenues for exploring hemoglobin-mediated stress responses in nitrogen-fixing tissues.

Together, these findings provide new insights into the molecular regulation of nitrogen fixation and nodule senescence in soybean. The identification of miR169-*NF-YA* highlights potential regulatory links between nitrogen signaling and nodule function. The discovery of miR2111 in nodules, alongside *TML* transcripts, suggests a putative miR2111-*TML* pathway that may locally modulate nitrogen fixation, extending beyond its established role in systemic AON. The transcriptional shifts in *Bradyrhizobium* during senescence, particularly the activation of the *paa* operon and stress-associated *rpoH*, reinforce a bacterial transition toward a free-living state as host support declines. Finally, the identification of *LOX* genes as potential nitrogen fixation biomarkers and the discovery of *Hb2* in nodules expand our understanding of oxygen and lipid metabolism in symbiosis, providing new avenues for investigating their roles in symbiotic efficiency. These findings underscore the complexity of soybean-*Bradyrhizobium* symbiosis and highlight promising regulatory targets for improving biological nitrogen fixation.

## MATERIALS AND METHODS

### Plant and Bacteria Growth

All plants were cultivated in pots within the greenhouse facilities at the Donald Danforth Plant Science Center (St. Louis, Missouri, USA). They were subjected to a 14/10 hour light/dark cycle and maintained at temperatures of 25/20°C, respectively. Soybean seeds (*Glycine max cv.* Williams-82) were sown into 1-gallon pots (two seeds per pot) containing a pre-rinsed substrate, consisting of a three to one ratio of vermiculite:perlite (3:1). Plants were well-watered and fertilized regularly via Dosatron with *reverse osmosis* water and B&D nutrient solution for growing legumes (with the addition of minimal nitrogen - 0.5 mM KNO_3_) (Broughton and Dilworth 1971). The lesser of the two plantlets was sacrificed following the seven-day germination period. Seven-day-old seedlings were then inoculated with 5ml of *Bradyrhizobium diazoefficiens* USDA110 diluted to 0.05 OD_600nm_. *Bradyrhizobium* (USDA-ARS culture collection, NRRL B-4361) was grown aerobically at 28°C in HM liquid medium at 6.8 pH prior to inoculation (Tóth et al. 2016).

### Acetylene Reduction Assay (ARA) and calculation of Nitrogenase Activity

Nitrogenase activity was measured with the acetylene reduction assay, as detailed by (Montes-Luz et al. 2023). ARA measurements began at 4 wpi and continued weekly until 7 wpi, typically occurring between 10 and 11 am. Each plant was carefully excavated from its soil substrate, with a 10-minute interval between each extraction to ensure continuity between downstream gas chromatography with flame ionized detection (GC-FID) injections. Immediately following extraction, roots and shoots were separated and weighed independently. The entire root system was then briefly rinsed, dried, and placed in a 1 L airtight amber, wide-mouth, glass bottle with septum inlet (SKU 174832 Sci. Spec. Service Inc.). Using a large syringe, we exchanged 5% of the air (50 mL) with acetylene gas, which was prepared in-house via the calcium carbide gas reaction, and incubated for 30 minutes at room temperature. Following incubation, a 500 µL gas sample was removed from the bottle and injected into a ThermoFisher Trace GC Ultra GC-FID; column #CP7584 PoraPLOT U (25 m x 0.53 mm x 20 um). Gas samples were extracted and injected at two-minute intervals. A total of five gas injections were performed for each biological replicate over a 10-minute period, from 30 to 40 minutes after the start of the initial acetylene incubation period.

Immediately after ARA sampling, all root nodules were harvested from their respective root systems and weighed three hours after the start of the incubation period. Nodules were then hand counted. Each root system served as a single biological replicate. Five root systems were sampled weekly. Four of the five plants underwent incubation with acetylene gas, while the fifth served as a negative gas control and remained untreated. Nitrogenase activity was calculated as the rate of ethylene produced―in nmols―per mg of nodule tissue per hour. An ethylene standard curve was generated and used to determine the amount of ethylene in each injection.

### Sampling (Nodule Collection)

Separate from plants designated for ARA sampling, soybean root nodules, aged 2 to 10 wpi, were harvested weekly, typically between 9 and 10am. Nodules were carefully collected in 2 ml screw cap tubes, immersed in liquid nitrogen, and promptly stored at −80°C. To ensure sample consistency, each technical replicate consisted of three nodules collected in close proximity to the root crown of a single plant. One biological replicate corresponds to one plant. From each biological replicate, we collected four technical replicates to facilitate RNA extraction.

### RNA Extraction

Frozen nodules were manually ground using pre-chilled (using liquid N_2_) mortar and pestles. Ground tissue was further pulverized using a MM400 tissuelyzer (Retsch, Germany): 2x 45sec at 30Hz. Total nodule RNA was extracted using TRIreagent (Ambion, USA), following manufacturer’s instructions plus an additional isopropanol rinse to remove remaining polysaccharides.

### RNA and small RNA sequencing and analysis

For RNA sequencing, we used 1 ug of DNAse I-treated total RNA (ThermoFisher Scientific, USA) for bacterial rRNA depletion using the commercial RiboMinus™ Bacteria 2.0 Transcriptome Isolation Kit (ThermoFisher Scientific, USA), directly followed by a second rRNA depletion using the plant specific commercial RiboMinus™ Plant Kit for RNA-Seq (ThermoFisher Scientific, USA). We then used the NEBNext® Ultra™ II Directional RNA Library Prep Kit for Illumina (New England Biolabs, USA) to generate libraries. To ensure correct library size capture, we performed a Bioanalyzer dsDNA HS chip assay on the Agilent 2100 Bioanalyzer (Agilent Technologies, Inc.) for each library. All processes followed manufacturers’ instructions. All libraries were sequenced on a NovaSeq2000 instrument using 100-bp paired-end reads.

For small RNA sequencing, we used 100 ng of DNAse I-treated total RNA (ThermoFisher Scientific, USA) as input for the RealSeq-AC version two kit (Realseq Biosciences, USA). To ensure correct library size capture, we performed a Bioanalyzer dsDNA HS chip assay on the Agilent 2100 Bioanalyzer (Agilent Technologies, Inc.) for each library. All libraries were sequenced on a NovaSeq2000 instrument using 50-bp single-end reads.

For small RNA data analysis, we trimmed the adaptors using Cutadapt version 1.16 (Martin 2011) using a minimum insert size of 10 nt. We assessed sequence quality using FastQC (http://www.bioinformatics.babraham.ac.uk/projects/fastqc/). We aligned clean reads to the soybean genome version *Gmax_*W82 v4.a1, from Soybase (Grant et al. 2010), and the *Bradyrhizobium* USDA110_ASM164267v1 genome from NCBI (NCBI Resource Coordinators et al. 2018). All subsequent analyses were performed using the software Bowtie2 (Langmead and Salzberg 2012). For miRNAs, we used ShortStack v4.4 and the latest version of miRBase (version 22; (Kozomara et al. 2019) to perform a *de novo* annotation of potential miRNAs that may have been previously overlooked. ShortStack identified 180 confident miRNAs, and 196 miRNA candidates, all of which were already annotated in miRBase v22. Using the recommended parameters, we did not detect any novel miRNAs, so we proceeded with the annotated soybean miRNAs from miRBase for subsequent analyses. Of the 756 annotated miRNAs in soybean, we successfully identified 748, with 581 exhibiting at least five reads at one or more time points (Supplemental Table 4).

For the RNA-seq libraries were analyzed using HiSat2 and Stringtie pipeline (Pertea et al. 2016), and the annotation files of the genomes mentioned before. We performed differential accumulation analyses using DESeq2 with default parameters, using reads that were not normalized as input (Love et al. 2014). In DESeq2, p-values were calculated using the Wald test and corrected for multiple testing using the Benjamini and Hochberg procedure. We generated graphical representations using the software ggplot2 (Wickham 2016) in the R statistical environment. Correlation analysis were performed using Spearman’s test at *p* < 0.05 and the corrplot R package (R Package “Corrplot” Visualization of a Correlation Matrix. - References - Scientific Research Publishing 2021).

### GO-term Enrichment (Shiny GO V0.80)

We used *Shiny GO V0.80* software (Ge et al. 2020) for Gene Ontology (GO) enrichment analysis of DE and hierarchical clustered soybean genes, with the recommended parameters and a False Discovery Rate cutoff 0.01. For GO characterization without enrichment, we used the custom-made software of Soybase (Grant et al. 2010). In both cases, the *Gmax*_v4.a1 genome and annotation was used as a reference.

## Acknowledgements

We thank Doug Allen of the Donald Danforth Plant Science Center (St. Louis, Missouri, USA) for providing soybean seed stocks used in this research and Gary Stacey of the University of Missouri (Columbia, Missouri, USA) for gifting the *Bradyrhizobium* strain. We are grateful to Bruna Montes Luz (University of Missouri, Columbia, Missouri, USA) for training in acetylene reduction assays (ARA) and to Anindita Banerjee and Michelle Liberton of the Himadri Pakrasi Lab at Washington University (St. Louis, Missouri, USA) for access to their lab and equipment for ARA practice and sample optimization. For all experimental ARA analyses, we used the GC-FID equipment in Doug Allen’s lab. Special thanks to Georgia Teague for her assistance in generating seed stocks and to Joanna Friesner for her invaluable contributions in editing this manuscript.

## Author Contributions

R.D. conceptualized the study with input from P.B., cultivated and inoculated soybeans, and harvested nodule samples. M.M. generated RNA and sRNA libraries. R.D. and S.M. conducted acetylene reduction assays. R.D. analyzed the data with input from P.B. and wrote the manuscript with contributions and interpretation from all authors.

## Current address of R.DelPercio

Blake Meyers Lab, Genome Center, University of California, Davis, CA, 95616

## Data availability

All RNA sequencing data has been deposited into the Gene Expression Omnibus under the accession code GSE289665

## REFERENCES

Anderson, C. R., Jensen, E. O., LLewellyn, D. J., Dennis, E. S., and Peacock, W. J. 1996. A new hemoglobin gene from soybean: a role for hemoglobin in all plants. Proc. Natl. Acad. Sci. U. S. A. 93:5682–5687

Andreou, A., and Feussner, I. 2009. Lipoxygenases - Structure and reaction mechanism. Phytochemistry. 70:1504–1510

Bergersen, F. J. 1970. The quantitative relationship between nitrogen fixation and the acetylene-reduction assay. Aust. J. Biol. Sci. 23:1015

Borges, F., and Martienssen, R. A. 2015. The expanding world of small RNAs in plants. Nat. Rev. Mol. Cell Biol. 16:727–741

Broughton, W. J., and Dilworth, M. J. 1971. Control of leghaemoglobin synthesis in snake beans. Biochem. J. 125:1075–1080

Cafiero, J. H., Salvetti Casasco, M., Lozano, M. J., Vacca, C., López García, S. L., Draghi, W. O., Lagares, A., and Del Papa, M. F. 2023. Genomic analysis of Sinorhizobium meliloti LPU63, an acid-tolerant and symbiotically efficient alfalfa-nodulating rhizobia. Front. Agron. 5:1175524

Cai, Z., Wang, Y., Zhu, L., Tian, Y., Chen, L., Sun, Z., Ullah, I., and Li, X. 2017. GmTIR1/GmAFB3-based auxin perception regulated by miR393 modulates soybean nodulation. New Phytol. 215:672–686

Cervantes-Pérez, S. A., Zogli, P., Amini, S., Thibivilliers, S., Tennant, S., Hossain, M. S., Xu, H., Meyer, I., Nooka, A., Ma, P., Yao, Q., Naldrett, M. J., Farmer, A., Martin, O., Bhattacharya, S., Kläver, J., and Libault, M. 2024. Single-cell transcriptome atlases of soybean root and mature nodule reveal new regulatory programs that control the nodulation process. Plant Commun. 5:100984

Chakraborty, S., Valdés-López, O., Stonoha-Arther, C., and Ané, J.-M. 2022. Transcription Factors Controlling the Rhizobium–Legume Symbiosis: Integrating Infection, Organogenesis and the Abiotic Environment. Plant Cell Physiol. 63:1326–1343

Chaulagain, D., Schnabel, E., Kappes, M., Lin, E. X., Müller, L. M., and Frugoli, J. A. 2023. TML1 AND TML2 SYNERGISTICALLY REGULATE NODULATION AND AFFECT ARBUSCULAR MYCORRHIZA IN MEDICAGO TRUNCATULA. Molecular Biology.: 2023.12.07.570674

Chen, H.-M., Chen, L.-T., Patel, K., Li, Y.-H., Baulcombe, D. C., and Wu, S.-H. 2010. 22-Nucleotide RNAs trigger secondary siRNA biogenesis in plants. Proc. Natl. Acad. Sci. U. S. A. 107:15269–15274

Ciampitti, I. A., de Borja Reis, A. F., Córdova, S. C., Castellano, M. J., Archontoulis, S. V., Correndo, A. A., Antunes De Almeida, L. F., and Moro Rosso, L. H. 2021. Revisiting biological nitrogen fixation dynamics in soybeans. Front. Plant Sci. 12:727021

Cuperus, J. T., Fahlgren, N., and Carrington, J. C. 2011. Evolution and functional diversification of MIRNA genes. Plant Cell. 23:431–442

De Luis, A., Markmann, K., Cognat, V., Holt, D. B., Charpentier, M., Parniske, M., Stougaard, J., and Voinnet, O. 2012. Two microRNAs linked to nodule infection and nitrogen-fixing ability in the legume Lotus japonicus. Plant Physiol. 160:2137–2154

Duan, L., Li, S.-J., Su, C., Sirichamorn, Y., Han, L.-N., Ye, W., Lôc, P. K., Wen, J., Compton, J. A., Schrire, B., Nie, Z.-L., and Chen, H.-F. 2021. Phylogenomic framework of the IRLC legumes (Leguminosae subfamily Papilionoideae) and intercontinental biogeography of tribe Wisterieae. Mol. Phylogenet. Evol. 163:107235

Du, M., Gao, Z., Li, X., and Liao, H. 2020. Excess nitrate induces nodule greening and reduces transcript and protein expression levels of soybean leghaemoglobins. Ann. Bot. 126:61–72

Dupont, L., Alloing, G., Pierre, O., El Msehli, S., Hopkins, J., Hérouart, D., and Frendo, P. 2012. The legume root nodule: From symbiotic nitrogen fixation to senescence.

Fan, K., Wong-Bajracharya, J., Lin, X., Ni, M., Ku, Y.-S., Li, M.-W., Tian, C. F., Chan, T.-F., and Lam, H.-M. 2021. Differentially expressed microRNAs that target functional genes in mature soybean nodules. Plant Genome. 14:e20103

Ferguson, B. J., Mens, C., Hastwell, A. H., Zhang, M., Su, H., Jones, C. H., Chu, X., and Gresshoff, P. M. 2019. Legume nodulation: The host controls the party. Plant Cell Environ. 42:41–51

Franck, S., Strodtman, K. N., Qiu, J., and Emerich, D. W. 2018. Transcriptomic Characterization of Bradyrhizobium diazoefficiens Bacteroids Reveals a Post-Symbiotic, Hemibiotrophic-Like Lifestyle of the Bacteria within Senescing Soybean Nodules. Int. J. Mol. Sci. 19

Fu, M., Yao, X., Li, X., Liu, J., Bai, M., Fang, Z., Gong, J., Guan, Y., and Xie, F. 2024. GmNLP1 and GmNLP4 activate nitrate-induced CLE peptides NIC1a/b to mediate nitrate-regulated root nodulation. Plant J.

Ge, S. X., Jung, D., and Yao, R. 2020. ShinyGO: a graphical gene-set enrichment tool for animals and plants. Bioinformatics. 36:2628–2629

Gordon, A. J., Minchin, F. R., James, C. L., and Komina, O. 1999. Sucrose synthase in legume nodules is essential for nitrogen fixation. Plant Physiol. 120:867–878

Grant, D., Nelson, R. T., Cannon, S. B., and Shoemaker, R. C. 2010. SoyBase, the USDA-ARS soybean genetics and genomics database. Nucleic Acids Res. 38:D843–6

Gresshoff, P. M., Su, H., Hastwell, A., Zhang, M., Grundy, E. B., Chu, X., and Ferguson, B. J. 2023. Functional genomics dissection of the nodulation autoregulation pathway (AON) in soybean (Glycine max). Research Square.

Guo, B., Zhang, J., Yang, C., Dong, L., Ye, H., Valliyodan, B., Nguyen, H. T., and Song, L. 2023. The late embryogenesis abundant proteins in soybean: Identification, expression analysis, and the roles of GmLEA4_19 in drought stress. Int. J. Mol. Sci. 24:14834

Gupta, K. J., Hebelstrup, K. H., Mur, L. A. J., and Igamberdiev, A. U. 2011. Plant hemoglobins: important players at the crossroads between oxygen and nitric oxide. FEBS Lett. 585:3843–3849

Haag, A. F., Arnold, M. F. F., Myka, K. K., Kerscher, B., Dall’Angelo, S., Zanda, M., Mergaert, P., and Ferguson, G. P. 2013. Molecular insights into bacteroid development during Rhizobium-legume symbiosis. FEMS Microbiol. Rev. 37:364–383

Hardy, R. W., Holsten, R. D., Jackson, E. K., and Burns, R. C. 1968. The acetylene-ethylene assay for n(2) fixation: laboratory and field evaluation. Plant Physiol. 43:1185–1207

Hill, R., Hargrove, M., and Arredondo-Peter, R. 2016. Phytoglobin: a novel nomenclature for plant globins accepted by the globin community at the 2014 XVIII conference on Oxygen-Binding and Sensing Proteins. F1000Res. 5:212

Hoang, N. T., Tóth, K., and Stacey, G. 2020. The role of microRNAs in the legume-Rhizobium nitrogen-fixing symbiosis. J. Exp. Bot. 71:1668–1680

Hooker, J. C., Smith, M., Zapata, G., Charette, M., Luckert, D., Mohr, R. M., Daba, K. A., Warkentin, T. D., Hadinezhad, M., Barlow, B., Hou, A., Lefebvre, F., Golshani, A., Cober, E. R., and Samanfar, B. 2023. Differential gene expression provides leads to environmentally regulated soybean seed protein content. Front. Plant Sci. 14:1260393

Huang, S., Zhou, J., Gao, L., and Tang, Y. 2021. Plant miR397 and its functions. Funct. Plant Biol. 48:361–370

Hu, Y., and Ribbe, M. W. 2013. Biosynthesis of the iron-molybdenum cofactor of nitrogenase. J. Biol. Chem. 288:13173–13177

Indrasumunar, A., Searle, I., Lin, M.-H., Kereszt, A., Men, A., Carroll, B. J., and Gresshoff, P. M. 2011. Nodulation factor receptor kinase 1α controls nodule organ number in soybean (Glycine max L. Merr). Plant J. 65:39–50

Iwakawa, H.-O., and Tomari, Y. 2013. Molecular insights into microRNA-mediated translational repression in plants. Mol. Cell. 52:591–601

Jiang, J., Zhu, H., Li, N., Batley, J., and Wang, Y. 2022. The miR393-Target Module Regulates Plant Development and Responses to Biotic and Abiotic Stresses. Int. J. Mol. Sci. 23

Jimenez-Vicente, E., Yang, Z.-Y., Ray, W. K., Echavarri-Erasun, C., Cash, V. L., Rubio, L. M., Seefeldt, L. C., and Dean, D. R. 2018. Sequential and differential interaction of assembly factors during nitrogenase MoFe protein maturation. J. Biol. Chem. 293:9812–9823

Kazmierczak, T., Yang, L., Boncompagni, E., Meilhoc, E., Frugier, F., Frendo, P., Bruand, C., Gruber, V., and Brouquisse, R. 2020. Chapter Seven - Legume nodule senescence: a coordinated death mechanism between bacteria and plant cells. Pages 181–212 in: Advances in Botanical Research, P. Frendo, F. Frugier, and C. Masson-Boivin, eds. Academic Press.

Kereszt, A., Mergaert, P., Montiel, J., Endre, G., and Kondorosi, É. 2018. Impact of Plant Peptides on Symbiotic Nodule Development and Functioning. Front. Plant Sci. 9:1026

Kikuchi, G., Motokawa, Y., Yoshida, T., and Hiraga, K. 2008. Glycine cleavage system: reaction mechanism, physiological significance, and hyperglycinemia. Proc. Jpn. Acad. Ser. B Phys. Biol. Sci. 84:246–263

Kim, Y., Wang, J., Ma, C., Jong, C., Jin, M., Cha, J., Wang, J., Peng, Y., Ni, H., Li, H., Yang, M., Chen, Q., and Xin, D. 2023. GmTCP and GmNLP Underlying Nodulation Character in Soybean Depending on Nitrogen. Int. J. Mol. Sci. 24

Koltun, A., Fuhrmann-Aoyagi, M. B., Cardoso Moraes, L. A., Lima Nepomuceno, A., Simões Azeredo Gonçalves, L., and Mertz-Henning, L. M. 2022. Uncovering the roles of hemoglobins in soybean facing water stress. Gene. 810:146055

Kosslak, R. M., Bookland, R., Barkei, J., Paaren, H. E., and Appelbaum, E. R. 1987. Induction of Bradyrhizobium japonicum common nod genes by isoflavones isolated from Glycine max. Proc. Natl. Acad. Sci. U. S. A. 84:7428–7432

Kozomara, A., Birgaoanu, M., and Griffiths-Jones, S. 2019. miRBase: from microRNA sequences to function. Nucleic Acids Res. 47:D155–D162

Kuzma, M. M., Hunt, S., and Layzell, D. B. 1993. Role of oxygen in the limitation and inhibition of nitrogenase activity and respiration rate in individual soybean nodules. Plant Physiol. 101:161–169

Kuźma, Winter, Storer, Oresnik, Atkins, and Layzell. 1999. The site of oxygen limitation in soybean nodules. Plant Physiol. 119:399–408

Laffont, C., Huault, E., Gautrat, P., Endre, G., Kalo, P., Bourion, V., Duc, G., and Frugier, F. 2019. Independent Regulation of Symbiotic Nodulation by the SUNN Negative and CRA2 Positive Systemic Pathways. Plant Physiol. 180:559–570

Langmead, B., and Salzberg, S. L. 2012. Fast gapped-read alignment with Bowtie 2. Nat. Methods. 9:357–359

Lepetit, M., and Brouquisse, R. 2023. Control of the rhizobium-legume symbiosis by the plant nitrogen demand is tightly integrated at the whole plant level and requires inter-organ systemic signaling. Front. Plant Sci. 14:1114840

Leyva-González, M. A., Ibarra-Laclette, E., Cruz-Ramírez, A., and Herrera-Estrella, L. 2012. Functional and transcriptome analysis reveals an acclimatization strategy for abiotic stress tolerance mediated by Arabidopsis NF-YA family members. PLoS One. 7:e48138

Li, J., Song, Q., Zuo, Z.-F., and Liu, L. 2022a. MicroRNA398: A Master Regulator of Plant Development and Stress Responses. Int. J. Mol. Sci. 23

Lim, C. W., Lee, Y. W., Lee, S. C., and Hwang, C. H. 2014. Nitrate inhibits soybean nodulation by regulating expression of CLE genes. Plant Sci. 229:1–9

Li, S., Lyu, X., Wang, X., Zhao, S., Ma, C., Yan, C., and Gong, Z. 2023. Assimilation of Nitrate into Asparagine for Transport in Soybeans. Agronomy. 13:2767

Li, Y., Pei, Y., Shen, Y., Zhang, R., Kang, M., Ma, Y., Li, D., and Chen, Y. 2022b. Progress in the Self-Regulation System in Legume Nodule Development-AON (Autoregulation of Nodulation). Int. J. Mol. Sci. 23

Lorio, J. C., Kim, W.-S., Krishnan, A. H., and Krishnan, H. B. 2010. Disruption of the Glycine cleavage system enables *Sinorhizobium fredii* USDA257 to form nitrogen-fixing nodules on agronomically improved North American soybean cultivars. Appl. Environ. Microbiol. 76:4185–4193

Love, M. I., Huber, W., and Anders, S. 2014. Moderated estimation of fold change and dispersion for RNA-seq data with DESeq2. Genome Biol. 15:550

Luan, M., Xu, M., Lu, Y., Zhang, Q., Zhang, L., Zhang, C., Fan, Y., Lang, Z., and Wang, L. 2014. Family-wide survey of miR169s and NF-YAs and their expression profiles response to abiotic stress in maize roots. PLoS One. 9:e91369

Martínez-Salazar, J. M., Sandoval-Calderón, M., Guo, X., Castillo-Ramírez, S., Reyes, A., Loza, M. G., Rivera, J., Alvarado-Affantranger, X., Sánchez, F., González, V., Dávila, G., and Ramírez-Romero, M. A. 2009. The Rhizobium etli RpoH1 and RpoH2 sigma factors are involved in different stress responses. Microbiology. 155:386–397

Martin, M. 2011. Cutadapt removes adapter sequences from high-throughput sequencing reads. EMBnet.journal. 17:10–12

Mergaert, P., Nikovics, K., Kelemen, Z., Maunoury, N., Vaubert, D., Kondorosi, A., and Kondorosi, E. 2003. A novel family in Medicago truncatula consisting of more than 300 nodule-specific genes coding for small, secreted polypeptides with conserved cysteine motifs. Plant Physiol. 132:161–173

Mergaert, P., Uchiumi, T., Alunni, B., Evanno, G., Cheron, A., Catrice, O., Mausset, A.-E., Barloy-Hubler, F., Galibert, F., Kondorosi, A., and Kondorosi, E. 2006. Eukaryotic control on bacterial cell cycle and differentiation in the Rhizobium-legume symbiosis. Proc. Natl. Acad. Sci. U. S. A. 103:5230–5235

Montes-Luz, B., Conrado, A. C., Ellingsen, J. K., Monteiro, R. A., de Souza, E. M., and Stacey, G. 2023.Acetylene Reduction Assay: A Measure of Nitrogenase Activity in Plants and Bacteria. Curr Protoc. 3:e766

Morell, M., and Copeland, L. 1985. Sucrose synthase of soybean nodules. Plant Physiol. 78:149–154

Müller, J., Wiemken, A., and Boller, T. 2001. Redifferentiation of bacteria isolated from Lotus japonicus root nodules colonized by Rhizobium sp. NGR234. J. Exp. Bot. 52:2181–2186

NCBI Resource Coordinators, Agarwala, R., Barrett, T., Beck, J., Benson, D. A., Bollin, C., Bolton, E., Bourexis, D., Brister, J. R., Bryant, S. H., Canese, K., Cavanaugh, M., Charowhas, C., Clark, K., Dondoshansky, I., Feolo, M., Fitzpatrick, L., Funk, K., Geer, L. Y., Gorelenkov, V., Graeff, A., Hlavina, W., Holmes, B., Johnson, M., Kattman, B., Khotomlianski, V., Kimchi, A., Kimelman, M., Kimura, M., Kitts, P., Klimke, W., Kotliarov, A., Krasnov, S., Kuznetsov, A., Landrum, M. J., Landsman, D., Lathrop, S., Lee, J. M., Leubsdorf, C., Lu, Z., Madden, T. L., Marchler-Bauer, A., Malheiro, A., Meric, P., Karsch-Mizrachi, I., Mnev, A., Murphy, T., Orris, R., Ostell, J., O’Sullivan, C., Palanigobu, V., Panchenko, A. R., Phan, L., Pierov, B., Pruitt, K. D., Rodarmer, K., Sayers, E. W., Schneider, V., Schoch, C. L., Schuler, G. D., Sherry, S. T., Siyan, K., Soboleva, A., Soussov, V., Starchenko, G., Tatusova, T. A., Thibaud-Nissen, F., Todorov, K., Trawick, B. W., Vakatov, D., Ward, M., Yaschenko, E., Zasypkin, A., and Zbicz, K. 2018. Database resources of the National Center for Biotechnology Information. Nucleic Acids Res. 46:D8–D13

Nguyen, C. X., Dohnalkova, A., Hancock, C. N., Kirk, K. R., Stacey, G., and Stacey, M. G. 2023. Critical role for uricase and xanthine dehydrogenase in soybean nitrogen fixation and nodule development. Plant Genome. 16:e20171

Ni, Z., Hu, Z., Jiang, Q., and Zhang, H. 2013. GmNFYA3, a target gene of miR169, is a positive regulator of plant tolerance to drought stress. Plant Mol. Biol. 82:113–129

Ohlendorf, D. H., Lipscomb, J. D., and Weber, P. C. 1988. Structure and assembly of protocatechuate 3,4-dioxygenase. Nature. 336:403–405

Okuma, N., and Kawaguchi, M. 2021. Systemic Optimization of Legume Nodulation: A Shoot-Derived Regulator, miR2111. Front. Plant Sci. 12:682486

Oono, R., Schmitt, I., Sprent, J. I., and Denison, R. F. 2010. Multiple evolutionary origins of legume traits leading to extreme rhizobial differentiation. New Phytol. 187:508–520

Pertea, M., Kim, D., Pertea, G. M., Leek, J. T., and Salzberg, S. L. 2016. Transcript-level expression analysis of RNA-seq experiments with HISAT, StringTie and Ballgown. Nat. Protoc. 11:1650–1667

Price, M. N., Deutschbauer, A. M., and Arkin, A. P. 2021. Four families of folate-independent methionine synthases. PLoS Genet. 17:e1009342

Puppo, A., Groten, K., Bastian, F., Carzaniga, R., Soussi, M., Lucas, M. M., de Felipe, M. R., Harrison, J., Vanacker, H., and Foyer, C. H. 2005. Legume nodule senescence: roles for redox and hormone signalling in the orchestration of the natural aging process. New Phytol. 165:683–701

Reid, D. E., Ferguson, B. J., and Gresshoff, P. M. 2011. Inoculation- and nitrate-induced CLE peptides of soybean control NARK-dependent nodule formation. Mol. Plant. Microbe. Interact. 24:606–618

Reyero-Saavedra, M. D. R., Qiao, Z., Sánchez-Correa, M. D. S., Díaz-Pineda, M. E., Reyes, J. L., Covarrubias, A. A., Libault, M., and Valdés-López, O. 2017. Gene Silencing of Argonaute5 Negatively Affects the Establishment of the Legume-Rhizobia Symbiosis. Genes. 8

R Package “Corrplot” Visualization of a Correlation Matrix. - References - Scientific Research Publishing. 2021. Available at: https://www.scirp.org/reference/referencespapers?referenceid=3377798 [Accessed March 18, 2025].

Sandhu, A. K., Brown, M. R., Subramanian, S., and Brözel, V. S. 2023. Bradyrhizobium diazoefficiens USDA 110 displays plasticity in the attachment phenotype when grown in different soybean root exudate compounds. Front. Microbiol. 14:1190396

Schwember, A. R., Schulze, J., Del Pozo, A., and Cabeza, R. A. 2019. Regulation of Symbiotic Nitrogen Fixation in Legume Root Nodules. Plants. 8

Shiraku, M. L., Magwanga, R. O., Zhang, Y., Hou, Y., Kirungu, J. N., Mehari, T. G., Xu, Y., Wang, Y., Wang, K., Cai, X., Zhou, Z., and Liu, F. 2022. Late embryogenesis abundant gene LEA3 (Gh_A08G0694) enhances drought and salt stress tolerance in cotton. Int. J. Biol. Macromol. 207:700–714

Simon, S. A., Meyers, B. C., and Sherrier, D. J. 2009. MicroRNAs in the rhizobia legume symbiosis. Plant Physiol. 151:1002–1008

Singh, P., Arif, Y., Miszczuk, E., Bajguz, A., and Hayat, S. 2022. Specific roles of lipoxygenases in development and responses to stress in plants. Plants. 11:979

Singh, S., and Varma, A. 2017. Structure, Function, and Estimation of Leghemoglobin. Pages 309–330 in: Soil Biology, Springer International Publishing, Cham.

Smagghe, B. J., Hoy, J. A., Percifield, R., Kundu, S., Hargrove, M. S., Sarath, G., Hilbert, J.-L., Watts, R. A., Dennis, E. S., Peacock, W. J., Dewilde, S., Moens, L., Blouin, G. C., Olson, J. S., and Appleby, C. A. 2009. Review: correlations between oxygen affinity and sequence classifications of plant hemoglobins. Biopolymers. 91:1083–1096

Stacey, G., Libault, M., Brechenmacher, L., Wan, J., and May, G. D. 2006. Genetics and functional genomics of legume nodulation. Curr. Opin. Plant Biol. 9:110–121

Stein, O., and Granot, D. 2019. An overview of sucrose synthases in plants. Front. Plant Sci. 10:95

Streit, W. R., Schmitz, R. A., Perret, X., Staehelin, C., Deakin, W. J., Raasch, C., Liesegang, H., and Broughton, W. J. 2004. An evolutionary hot spot: the pNGR234b replicon of Rhizobium sp. strain NGR234. J. Bacteriol. 186:535–542

Tóth, K., Batek, J., and Stacey, G. 2016. Generation of Soybean (Glycine max) Transient Transgenic Roots. Curr Protoc Plant Biol. 1:1–13

Tsikou, D., Yan, Z., Holt, D. B., Abel, N. B., Reid, D. E., Madsen, L. H., Bhasin, H., Sexauer, M., Stougaard, J., and Markmann, K. 2018. Systemic control of legume susceptibility to rhizobial infection by a mobile microRNA. Science. 362:233–236

Udvardi, M., and Poole, P. S. 2013. Transport and metabolism in legume-rhizobia symbioses. Annu. Rev. Plant Biol. 64:781–805

Van de Velde, W., Guerra, J. C. P., De Keyser, A., De Rycke, R., Rombauts, S., Maunoury, N., Mergaert, P., Kondorosi, E., Holsters, M., and Goormachtig, S. 2006. Aging in legume symbiosis. A molecular view on nodule senescence in Medicago truncatula. Plant Physiol. 141:711–720

Vessey, J. 2004. Measurement of nitrogenase activity in legume root nodules: In defense of the acetylene reduction assay. Plant Soil. 158:151–162

Wang, C., Saldanha, M., Sheng, X., Shelswell, K. J., Walsh, K. T., Sobral, B. W. S., and Charles, T. C. 2007. Roles of poly-3-hydroxybutyrate (PHB) and glycogen in symbiosis of Sinorhizobium meliloti with Medicago sp. Microbiology. 153:388–398

Wang, H., and Gunsalus, R. P. 2003. Coordinate regulation of the Escherichia coli formate dehydrogenase fdnGHI and fdhF genes in response to nitrate, nitrite, and formate: roles for NarL and NarP. J. Bacteriol. 185:5076–5085

Wang, L., Sun, Z., Su, C., Wang, Y., Yan, Q., Chen, J., Ott, T., and Li, X. 2019. A GmNINa-miR172c-NNC1 Regulatory Network Coordinates the Nodulation and Autoregulation of Nodulation Pathways in Soybean. Mol. Plant. 12:1211–1226

Wang, Z., Wang, L., Wang, Y., and Li, X. 2020. The NMN module conducts nodule number orchestra. iScience. 23:100825

Wickham, H. 2016. ggplot2 - Elegant Graphics for Data Analysis. Springer International Publishing.

Xu, F., Liu, Q., Chen, L., Kuang, J., Walk, T., Wang, J., and Liao, H. 2013. Genome-wide identification of soybean microRNAs and their targets reveals their organ-specificity and responses to phosphate starvation. BMC Genomics. 14:66

Xu, H., Li, Y., Zhang, K., Li, M., Fu, S., Tian, Y., Qin, T., Li, X., Zhong, Y., and Liao, H. 2021. miR169c-NFYA-C-ENOD40 modulates nitrogen inhibitory effects in soybean nodulation. New Phytol. 229:3377–3392

Yan, Z., Hossain, M. S., Arikit, S., Valdés-López, O., Zhai, J., Wang, J., Libault, M., Ji, T., Qiu, L., Meyers, B. C., and Stacey, G. 2015. Identification of microRNAs and their mRNA targets during soybean nodule development: functional analysis of the role of miR393j-3p in soybean nodulation. New Phytol. 207:748–759

Yuan, S., Zhou, S., Feng, Y., Zhang, C., Huang, Y., Shan, Z., Chen, S., Guo, W., Yang, H., Yang, Z., Qiu, D., Chen, H., and Zhou, X. 2021. Identification of the Important Genes of Bradyrhizobium diazoefficiens 113-2 Involved in Soybean Nodule Development and Senescence. Front. Microbiol. 12:754837

Yun, J., Wang, C., Zhang, F., Chen, L., Sun, Z., Cai, Y., Luo, Y., Liao, J., Wang, Y., Cha, Y., Zhang, X., Ren, Y., Wu, J., Hasegawa, P. M., Tian, C., Su, H., Ferguson, B. J., Gresshoff, P. M., Hou, W., Han, T., and Li, X. 2023. A nitrogen fixing symbiosis-specific pathway required for legume flowering. Sci Adv. 9:eade1150

Zanetti, M. E., Rípodas, C., and Niebel, A. 2017. Plant NF-Y transcription factors: Key players in plant-microbe interactions, root development and adaptation to stress. Biochim. Biophys. Acta. 1860:645–654

Zhang, M., Su, H., Gresshoff, P. M., and Ferguson, B. J. 2021. Shoot-derived miR2111 controls legume root and nodule development. Plant Cell Environ. 44:1627–1641

Zhan, J., and Meyers, B. C. 2023. Plant Small RNAs: Their Biogenesis, Regulatory Roles, and Functions. Annu. Rev. Plant Biol. 74:21–51

Zhou, S., Zhang, C., Huang, Y., Chen, H., Yuan, S., and Zhou, X. 2021. Characteristics and Research Progress of Legume Nodule Senescence. Plants. 10

Zhu, F., Deng, J., Chen, H., Liu, P., Zheng, L., Ye, Q., Li, R., Brault, M., Wen, J., Frugier, F., Dong, J., and Wang, T. 2020. A CEP peptide receptor-like kinase regulates auxin biosynthesis and ethylene signaling to coordinate root growth and symbiotic nodulation in Medicago truncatula. Plant Cell. 32:2855–2877

Zougari, A., Guy, S., and Planchon, C. 1995. Genotypic lipoxygenase variation in soybean seeds and response to nitrogen nutrition. Plant Breed. 114:313–316

