## Supplemental Figures for "Transcriptional dynamics of nitrogen fixation and senescence in soybean nodules: A dual perspective on host and *Bradyrhizobium* regulation"

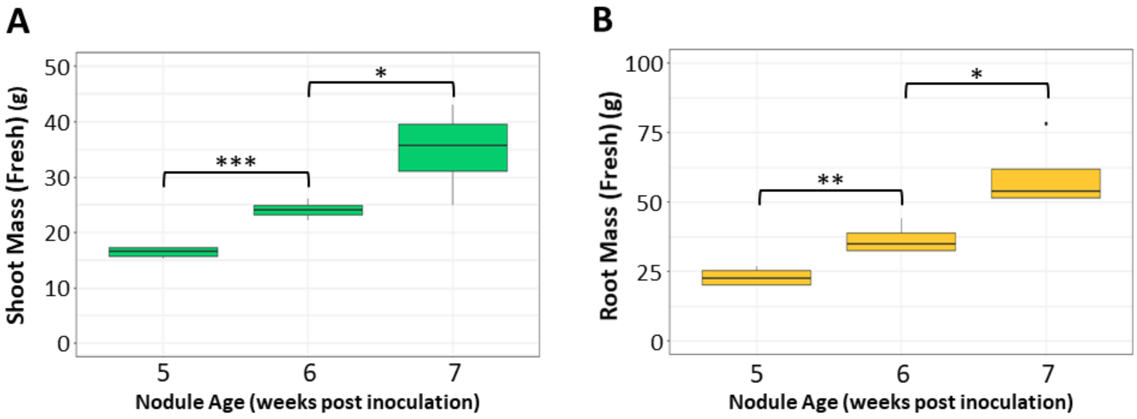


**Supplemental Figure 1. Shoot and root mass measured prior to Acetylene Reduction Assay (ARA)** Fresh weight of (A) shoot and (B) root of soybean plants used in the ARA. Data collected spans 5 to 7 weeks post inoculation (wpi) with *Bradyrhizobium* USDA110. Significant p-values are indicated by asterisks: * (0.01 to 0.05), ** (0.001 to 0.01), and *** (< 0.001). Note: No shoot and root mass measurements for 4 wpi samples.

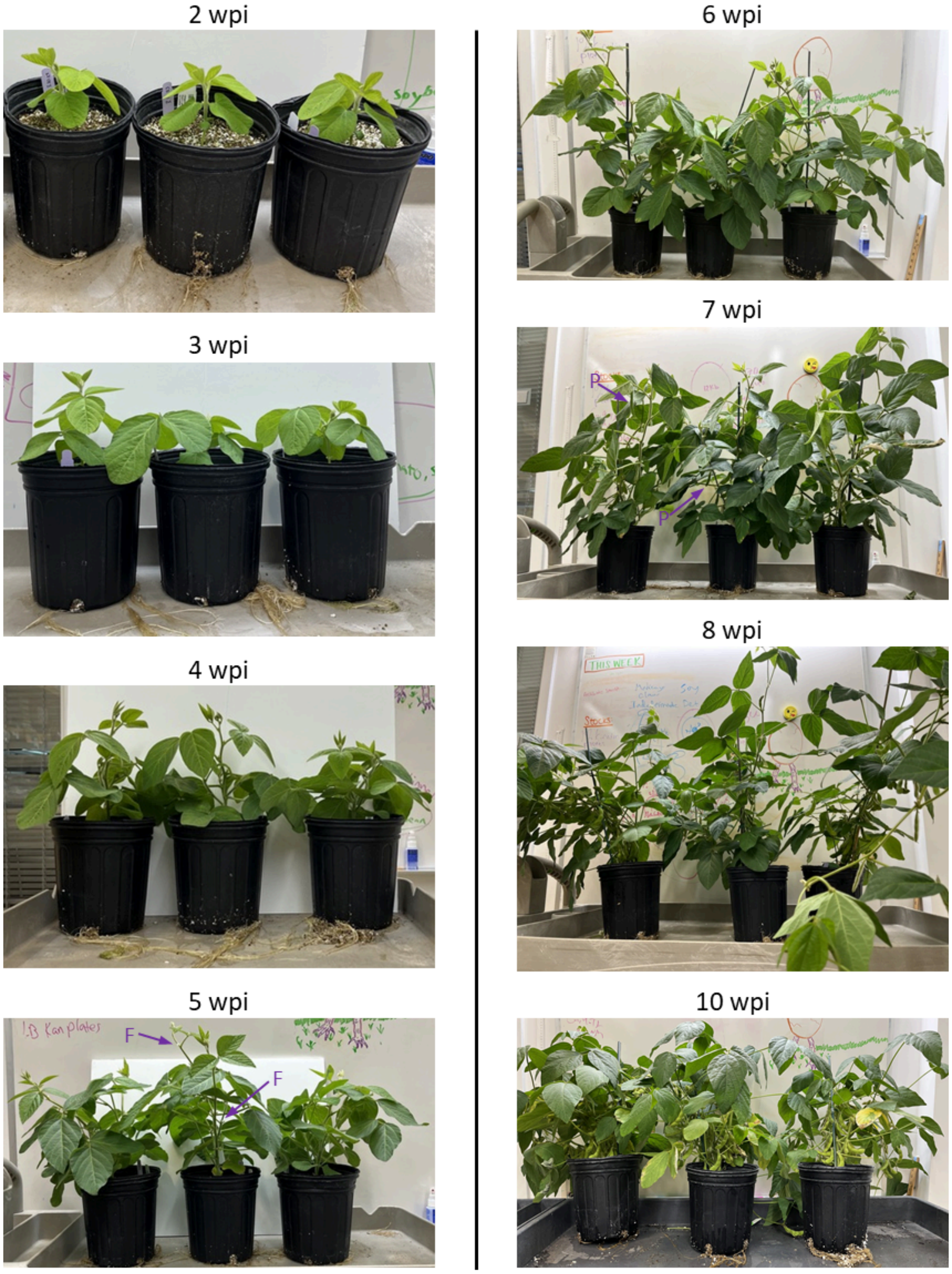


**Supplemental Figure 2. Developmental progression of soybean plants used for RNA and small RNA sequencing and metatranscriptomic analyses (2–10 weeks post inoculation)**
Representative images of soybean plants (*Glycine max cv.* Williams 82) grown for nodule collection and subsequent RNA sequencing (RNA-seq and small RNA-seq) to analyze both plant and *Bradyrhizobium* transcriptomes. Plants were sampled weekly from 2 to 10 wpi with *Bradyrhizobium diazoefficiens* USDA110. The images illustrate vegetative growth and the transition to reproductive stages, with **flowering (F)** observed beginning at 5 wpi and **pod development (P)** emerging around 7 wpi. These plants provided RNA for metatranscriptomic analysis of gene expression changes throughout early nodule development, nitrogen fixation, and senescence.

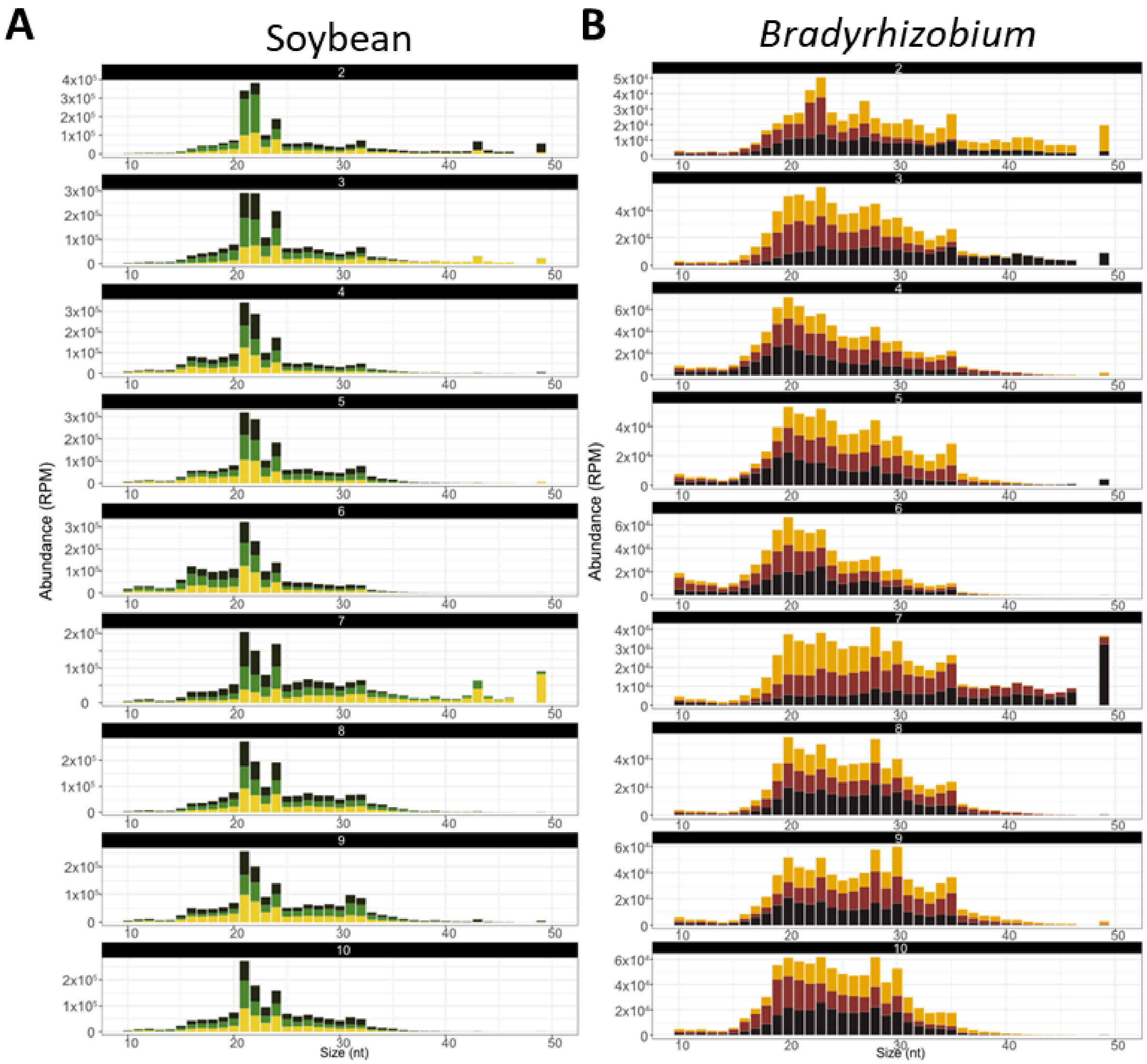


**Supplemental Figure 3. Distinct small RNA size distributions in soybean nodules reveal host and bacterial contributions**

(A) Read size distribution of small RNA (sRNA) reads mapping to the *Glycine max* (W82_v4.a1) genome, arranged from 2 to 10 weeks post-inoculation (wpi), top to bottom. The x-axis represents sRNA size (10–50 nucleotides), and the y-axis shows abundance in reads per million (RPM). Distinct peaks at 21, 22, and 24 nucleotides correspond to known plant miRNA and siRNA size classes, shifting from 22-nt dominance in early development to 21-nt prevalence post-4 wpi.

(B) Read size distribution of sRNA reads mapping to the *Bradyrhizobium diazoefficiens* USDA110 genome, arranged from 2 to 10 wpi. Unlike plant sRNAs, *Bradyrhizobium* sRNAs display a broader size range (17–35 nt) with no dominant peak, suggesting diverse bacterial regulatory mechanisms. Each color in the bar plots represents a different biological replicate, with three biological replicates per time point.

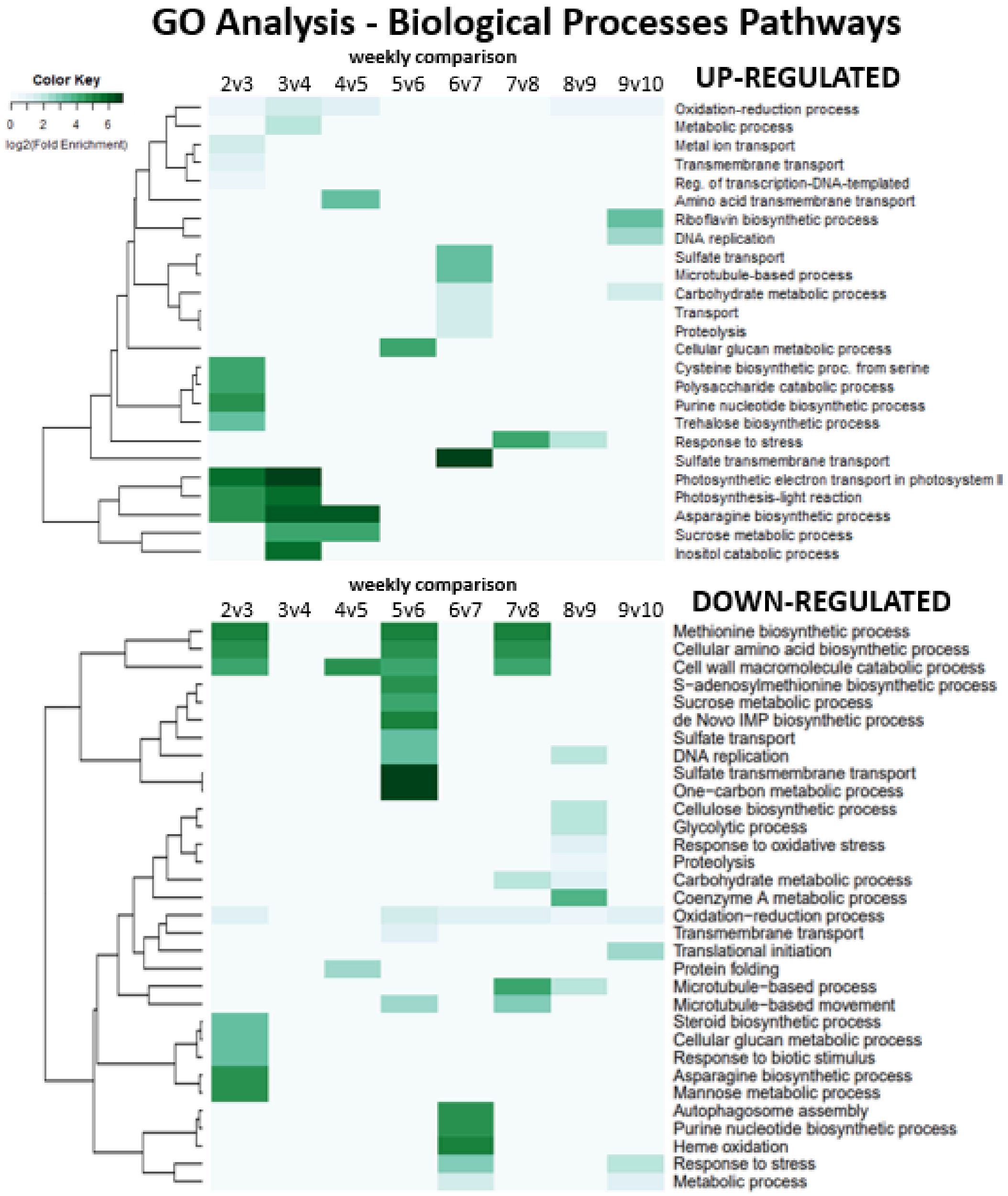


**Supplemental Figure 4. Temporal regulation of biological processes in soybean nodules identified through Gene Ontology (GO) analysis**

GO analysis of differentially expressed (DE) soybean genes reveals the temporal regulation of biological processes throughout nodule development. Upregulated (top) and downregulated (bottom) genes were analyzed using ShinyGO (v0.80) (Ge et al. 2020), with pathway enrichment scaled by the log2 of Fold Enrichment. DE gene expression data were derived from soybean nodules collected at weekly intervals from 2 to 10 weeks post-inoculation (wpi). Key biological processes associated with developmental stages include sucrose metabolic processes, asparagine biosynthesis, sulfate transport, and oxidation-reduction processes, providing insights into the dynamic transcriptional regulation underlying nodule function.

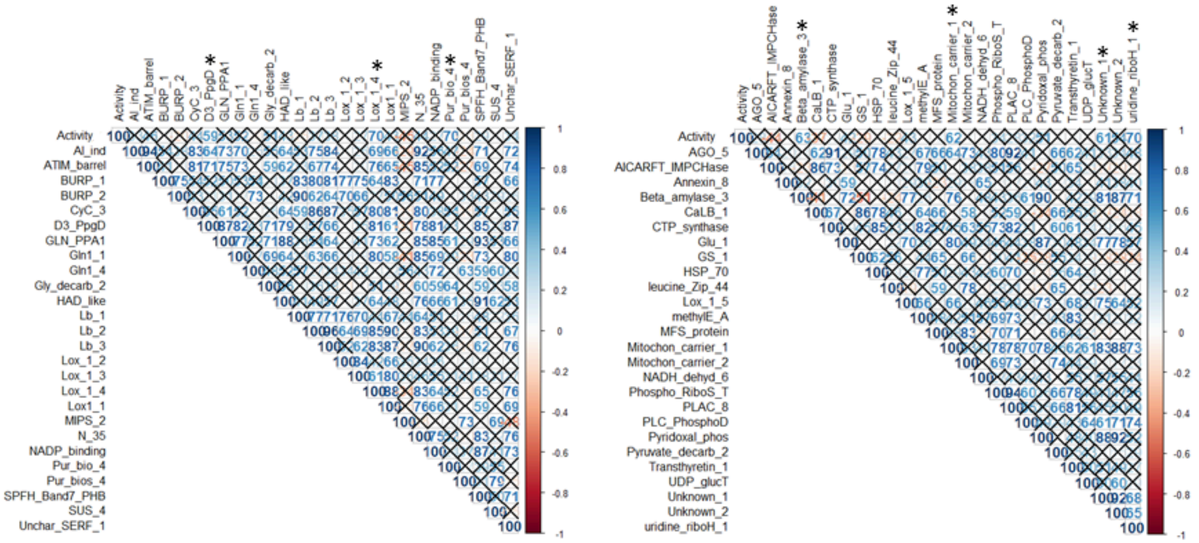


**Supplemental Figure 5. Correlation analysis of *2-up-5-down-10* differentially expressed soybean genes with nitrogenase activity**

Pearson correlation matrices of 52 highly expressed soybean *2-up-5-down-10* differentially expressed (DE) genes with nitrogenase activity. The left panel shows the 26 highest-expressing genes (2–10 wpi average FPKM), while the right panel shows the next 26 (27 to 52) highest-expressing *2-up-5-down-10* DE genes. Gene names were abbreviated for better visualization. Correlation values are displayed in blue, with darker shades indicating stronger correlations (only positive correlations were considered significant). Black crosses denote statistically insignificant correlations ($p > 0.05$). Asterisks (*) indicate genes significantly correlated with nitrogenase activity ($p < 0.05$), identified as candidate biomarkers for nitrogen fixation. Genes are sorted alphabetically within each correlation matrix.
